# Single Cell lineage Tracing Identifies Cancer Testis Antigens as Mediators of Chemoresistance in Small Cell Lung Cancer

**DOI:** 10.1101/2022.10.20.513051

**Authors:** Hannah Wollenzien, Yohannes Afeworki, Robert Szczepaniak-Sloane, Anthony Restaino, Michael S. Kareta

## Abstract

Small Cell Lung Cancer (SCLC) is often a heterogeneous tumor, where dynamic regulation of key transcription factors can drive multiple populations of phenotypically different cells which contribute differentially to tumor dynamics. This tumor is characterized by a very low 2-year survival rate, high rates of metastasis, and rapid acquisition of chemoresistance. The heterogeneous nature of this tumor makes it difficult to study and to treat, as it is not clear how or when this heterogeneity arises. Here we describe temporal, single-cell analysis of SCLC to investigate tumor initiation and chemoresistance in both SCLC xenografts and *in situ* SCLC mouse models. We identify an early population of tumor cells with high expression of AP-1 network genes that are critical for tumor growth. Furthermore, we have identified and validated the cancer testis antigens (CTAs) PAGE5 and GAGE2A as mediators of chemoresistance in human SCLC. CTAs have successfully been targeted in other tumor types and may be a promising avenue for targeted therapy in SCLC.

## Introduction

Small cell lung cancer (SCLC) is a devastating disease characterized by a 5-year survival rate of only 6% (1). The majority of patients (75%) present with extensive stage disease at diagnosis, and survival for these patients is generally under 1 year (2). Most patients are treated with first-line chemotherapy as the disease is commonly observed as extensive stage disease at diagnosis, and therefore surgical resection is rare (3). While initial response to chemotherapy generally appears favorable, the majority of patient’s tumors will rapidly acquire chemoresistance and relapse within months (2). Despite the therapeutic advances made in other tumor types, the grim outlook for patients with SCLC has been largely unchanged in the last 60 years (2, 4, 5). Many clinical trials have investigated the feasibility to utilize targeted therapies in SCLC, such as inhibition of PARP and other checkpoint inhibitors, inhibition of apoptotic pathways, Notch pathway inhibitors and others has shown either minimal responses or negative toxicities (6, 7). Recently, a PD-L1 inhibitor was approved for patients with extensive stage disease, but with an only two-month increase in overall survival time (8), further advances are critically needed.

SCLC is highly correlated with smoking, and almost universal loss of the tumor suppressors *RB* (*RB1*) and *p53* (*TP53*) is observed (9). SCLC historically has been thought of as a primarily neuroendocrine disease where patients can be classified in to “classic” or “variant” subtypes, with most patients being the neuroendocrine-high “classic” subtype (10). As our knowledge of SCLC genetics evolves, it has become increasingly clear that patients can be stratified in to one of four molecular subtypes based on key transcription factor expression: *NEUROD1*, *ASCL1*, *YAP1*, or *POU2F3* (11). These four molecular subtypes (SCLC-N, SCLC-A, SCLC-Y, SCLC-P, respectively) seem to have a differential effect on tumor behavior and patient outcome, but the regulation and behavior of subtypes is still largely unknown (11–16). Also unknown are the factors that lead to determining which subtype will be prominent, how dynamic an individual tumor’s subtype remains during tumor development, and how these factors relate to chemosensitivity/chemoresistance.

Cells in a single tumor are often phenotypically different from one another and therefore contribute differentially to tumor dynamics, a phenomenon known as intratumoral heterogeneity (ITH) (17–19). ITH is perhaps best demonstrated by the cancer stem cell model, which proposes a unique subpopulation of rare tumor cells that, despite their relative rarity as a population, are most directly responsible for tumor initiation, maintenance, and metastasis (20). ITH has an impact on response to therapy, particularly in SCLC, as demonstrated by patients’ almost full response to chemotherapy, only to have extensive disease re-occur rapidly (2, 21). This suggests either a population of cells inherently resistant to therapy or have the plasticity to adapt and become resistant when faced with therapeutic pressures. While ITH has been studied in many tumor types, in part driven by such resources as the TCGA, studies investigating ITH in SCLC are only slowly emerging (22).

ITH has been shown to occur in SCLC, particularly post-chemotherapy and metastatic events (13-15, 21, 23-28). As tumors progress, the predominant molecular subtype can change, which generally progresses from an *ASCL1*-high lineage to a *NEUROD1*-high lineage (14, 15). A number of potential regulators of SCLC identity, including *SOX2*, *MYC*, and *NOTCH* have emerged, however a definitive lineage has yet to be constructed (13–16, 21). A greater understanding of SCLC ITH is needed, knowing that ITH in SCLC poses a formidable challenge in providing successful treatments. To uncover or design better therapeutics for SCLC, it is first critical that we understand the contribution of different clonal populations to ITH, and how those populations change in response to therapy.

Historically, tumors have been sequenced using bulk RNA sequencing methods, however, this hides the true contribution of individual populations and aggregates the signal from the entire sample (23, 26). Single-cell RNA sequencing (scRNA-seq) has radically transformed our study and understanding of ITH. Indeed, scRNA-seq has generated important insights into SCLC tumors dynamics (14, 21, 28, 29), and combined with other genetic profiling has been used to generate pseudotime maps of ITH in response to growth and chemotherapy (13-15, 21, 23, 26, 30). While incredibly useful, pseudotime trajectories do not allow for the pinpointing of the populations critical for these tumor dynamics, rather phenotypic characteristics are inferred from the populations identified in the screening. In recent years, genetic barcode lineage tracing has emerged as a novel tool to trace individual clonal populations over time (31–33). Genetic barcode lineage tracing allows for the identification of individual cellular clones by inserting a unique piece of DNA to serve as a “barcode” in the genome of a cell using a retrovirus or CRISPR sgRNA library (28, 34, 35). As the cells divide, the barcode will be passed to the progeny, allowing for a high-resolution tracing of individual clonal populations. Originally used for tracing populations in hematopoiesis (31, 36), the use of genetic barcode lineage tracing to understand tumor heterogeneity has exploded in the last few years. It has now been used to understand cellular lineages and heterogeneity in a number of cancer types including glioblastoma, breast, non-small cell lung cancer, leukemia, and melanoma (32, 35, 37–45). Currently, genetic barcoding has been used to understand spatial architecture in SCLC xenografts and the role of PTEN in regulating these clonal structures (30), an expanded use of barcoding techniques in both SCLC mouse models and xenografts will be required to shed light on broader SCLC growth dynamics and chemoresistance. While the use of genetic barcoding to barcode tumors *in situ,* by the use of genetically engineered animal models of cancer is a newer advance on the genetic barcoding technology (28, 35), and its utility in SCLC is still emerging.

SCLC is almost always driven by loss of tumor suppressors *Rb* and *p53*, however, the very early stages of tumor development are not always captured in animal models that are aged out to three or even six months-post tumor initiation. We have yet to understand what happens in the very early stages after tumor initiation. Here we use genetic barcoding combined with scRNA-seq in a mouse model of SCLC and xenografts of both the SCLC-A and SCLC-N subtypes to understand tumor heterogeneity and evolution. We identify a population of tumor cells with high expression of AP-1 network genes that is maintained throughout the course of disease and is required for colony formation. In barcoded SCLC xenografts, the Cancer Testis Antigens *PAGE5* and *GAGE2A* were significantly upregulated in chemoresistant tumors and were validated to be mediators of chemoresistance in SCLC. Other members of the CTA family have been successfully targets for immunotherapy in other tumor types and could represent a promising therapeutic target for SCLC.

## Results

### Development of an *in situ* cellular barcoding model for SCLC

To barcode tumors as they form *in situ,* we used the well-characterized SCLC mouse model with Cre-mediated deletion of tumor suppressors *Rb, p53,* and *p130* (RPR2) (46), bred with an *H11^lox-stop-lox-Cas9^* mouse (47) to generate the RPR2-Cas9 mouse model (Fig. 1A). Upon intratracheal Cre injection, the mice rapidly develop tumors and concurrently express Cas9 (Supplementary Fig. S1). Mice were barcoded by intratracheal AAV injection at one-month intervals starting at time of Cre injection, through five months post-tumor initiation, and were euthanized at one-month intervals following barcoding (Fig. 1A, Supplementary Table S1). Designing the barcoding of lungs in this way allows for the barcoding of tumors at early, middle, and late stages of tumor development. After dissection, barcoded tumor cells were isolated via FACS, and used for scRNA-seq.

**Figure 1.**
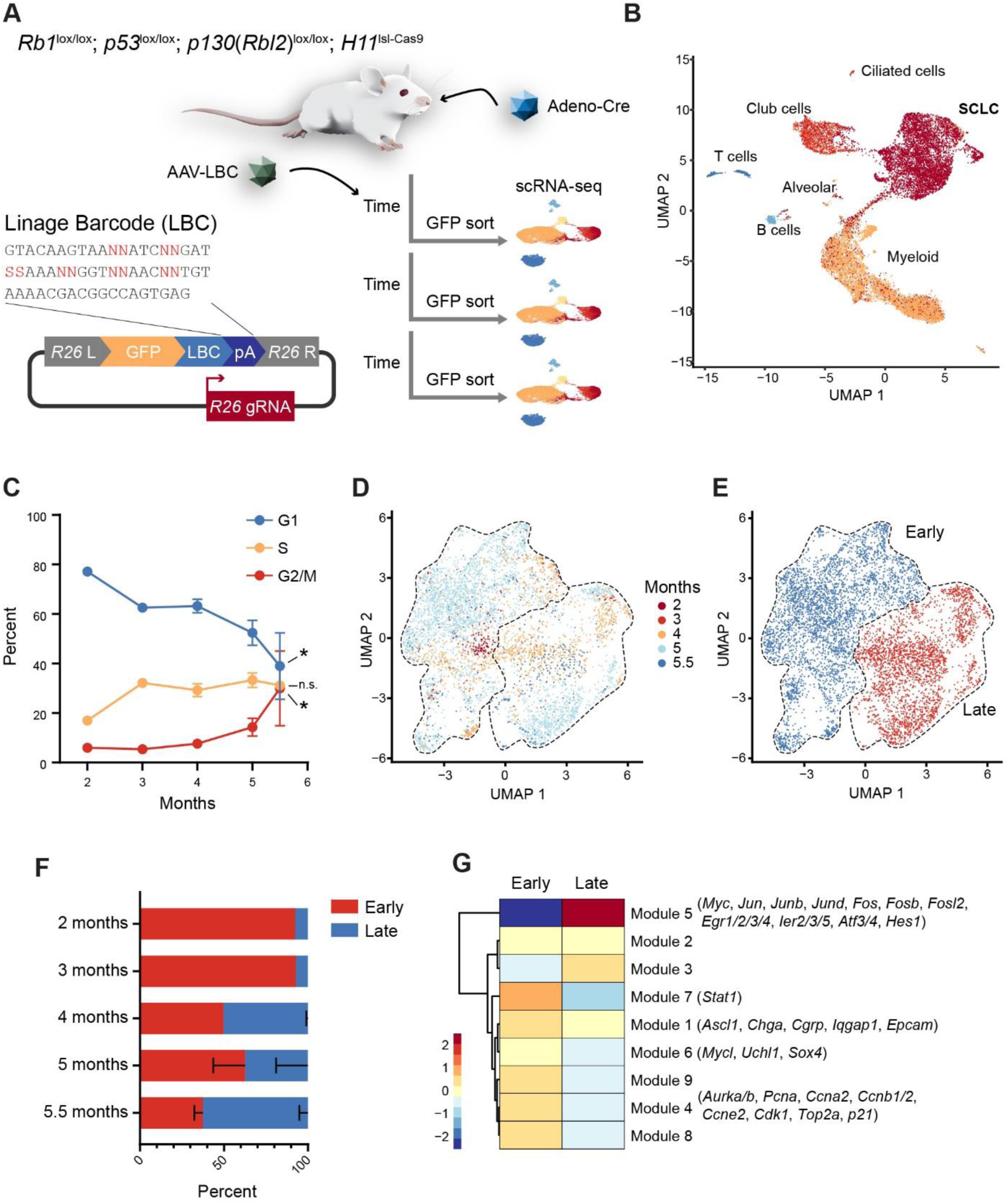
Single-Cell RNA sequencing reveals two distinct subpopulations in SCLC. **A,** Overview of the *in situ* barcoding model. Tumors in the RPR2-Cas9 model were initiated, and barcoded at one-month intervals, up to 5 months post-initiation. Tumors were harvested at one-month intervals following barcoding, up to 6 months, and GFP^+^ cells were isolated and scRNA-seq was performed. **B,** Multiple cellular populations were detected in the scRNA-seq due to the presence of microdissected tumors in this analysis, including a large population of SCLC cells. **C,** Proportion of cells in each stage of the cell cycle at varying months of tumor development. **D,** UMAP plot of the SCLC cells labeled by months of tumor development. Black outlines describe the clusters identified in (**E**). **E,** Unbiased clustering of the SCLC cells into two clusters. Due to the distribution of timepoints in (**D**), clusters are labeled “early” or “late”. **F,** Proportion of cells in the early and late populations at each timepoint after tumor initiation. **G,** Gene modules that differentiate early and late populations. Significance determined by an unpaired student’s t-test where (*) P<0.05, (**) P<0.01, (***) P<0.001.

### Single-cell RNA sequencing identifies two distinct signatures

SCLC tumor cells were readily identifiable form other cells of the tumor microenvironment by automatic annotation using known lung markers (Supplementary Fig. S2) and were characterized by classic SCLC neuroendocrine markers, *Ascl1*, and *Mycl* (Supplementary Fig. S3). Other cell types that are identified by this automatic annotation are myeloid, club, alveolar, ciliated cells, and T and B immune cells (Fig. 1B, Supplementary Fig. S2). Upon analysis of the scRNA-seq data, no barcodes were detectable in the tumors barcoded *in situ* (Supplementary Fig. S1). Despite having tumors that are immuno-reactive to antibodies against GFP, and the detection of GFP^+^ cells via flow cytometry, the depth of scRNA-seq in this case was not sufficient to pick up reads from the low-expressed GFP and barcode (Supplementary Fig. S4). Despite not detecting barcodes in these samples, due to the timewise design of the animal studies we then set out to determine what developmental pathways might be altered during SCLC development. When evaluating the cell cycle composition of the cells within the tumor populations, the proportion of cells in G1 decreases with each subsequent month that tumors are allowed to form, and the percentage of cells in G2 or metaphase increases nearly 6-fold after five months of tumor development (Fig. 1C, Supplementary Fig. S5). When plotting the distribution of tumor cells by developmental time, we observe an “early” tumor signature that arises in the tumors isolated after two months of development and persists through later stages of tumor development (Fig. 1D, 1E). In the later months of tumor development (month 4 and following), a “late” signature emerges, which eventually is responsible for the majority of the tumor (62.5% ±7.2) in the longest developed tumors (Fig. 1D - 1F). The early and late tumor populations can be distinguished by the gene expression of several distinct expression modules (Fig. 1G, Supplementary Table S3). We observe a decrease in neuroendocrine markers (including *Ascl1*, *Chga*, *Cgrp*, and *Mycl*) in the late tumor cells (modules 1 and 6) (14, 48), and regulation of many cell cycle genes which could drive the decrease of cells in G1 at later stages (module 4, Fig. 1C). Noteworthy however, was module 5 which is comprised of members of the AP-1 complex, and most strongly (P = 4.16 ×10^−7^) distinguishes between the early and late populations (Fig. 1G).

### The AP-1 network as a requirement for tumor maintenance and associated with chemoresistance

In evaluating the transcriptomic differences between the early and late tumor clusters, members of the AP-1 transcriptional network were not only upregulated in the late population but are also expressed in a majority of the cells indicating it is a potential driver of this group and not simply a rare subtype (Fig. 2A, 2B). Members of the AP-1 network including *Jun* and *Fos* are indeed implicated in tumor development (49–59) although their role has not yet been characterized in SCLC. We therefore sought to investigate the impact of disruption of the AP-1 complex on tumor development in SCLC. Using a dominant-negative *cJun* construct (JunDN) (60), AP-1 signaling was disrupted in the human SCLC cell lines NJH29 and NCI-H82, and cells were sorted by GFP expression from the JunDN vector to capture transduced cells. The cells were seeded into soft agar to assess colony formation abilities. SCLC cells with disruption of AP-1 signaling due to JunDN formed significantly fewer colonies (98.6% reduction, P = 5.0 x10^−5^) than wild-type cells (Fig. 2C, 2D) while cell viability was not affected (Fig. 2E). Two of the mice in this study were treated with clinically relevant doses of cisplatin and etoposide and allowed to progress until they reached ethical tumor endpoints. The majority (64.2%) of the chemoresistant cells from these tumors had gene expression signatures consistent with the late cell cluster, which has enrichment of AP-1 family members (Fig. 2F, 2G).

**Figure 2.**
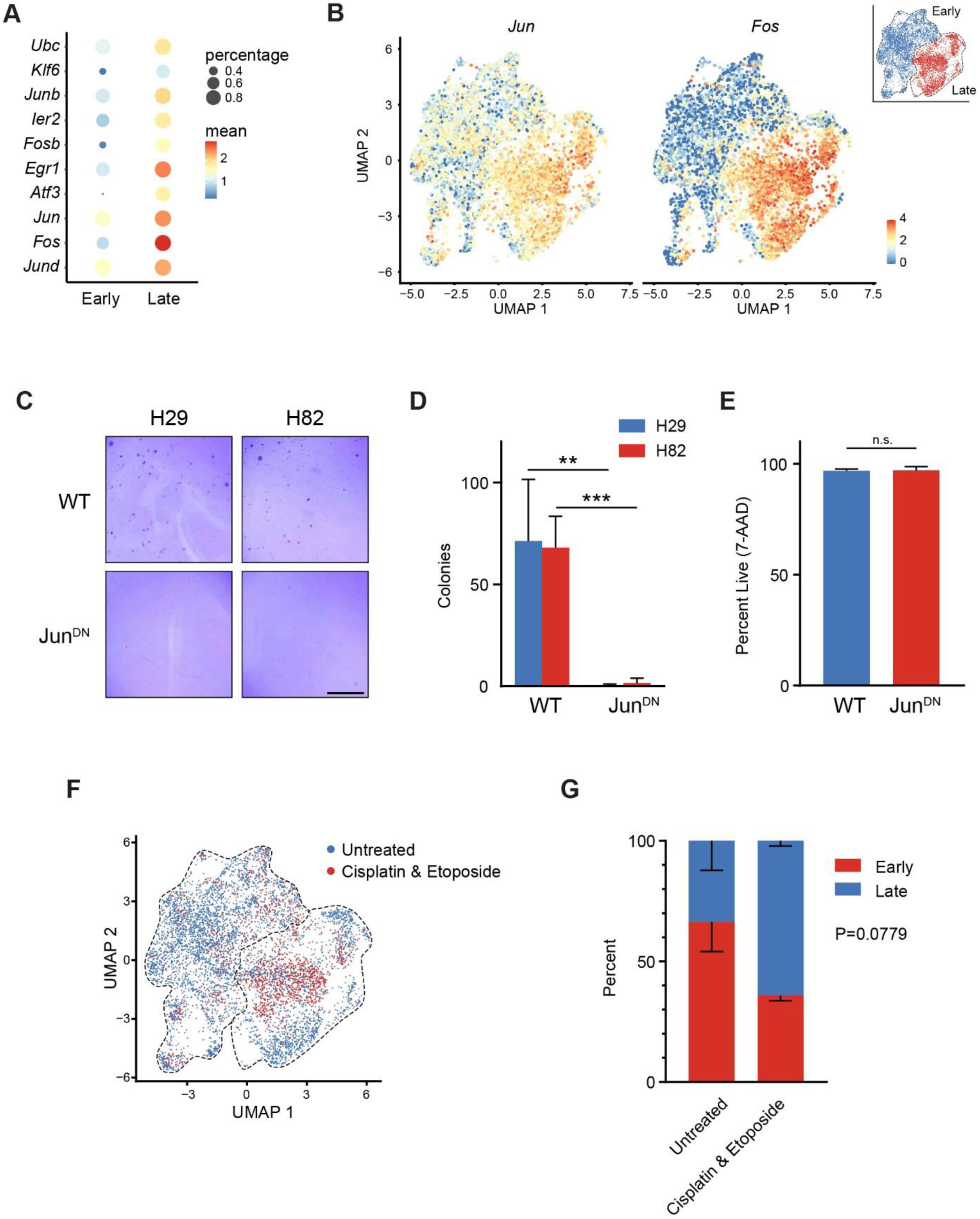
The AP-1 Network is required for tumor initiation and chemo-resistance. **A,** Members of the AP-1 network are upregulated in SCLC, and particularly in the Late cluster of cells. **B,** UMAP of *Fos* and *Jun* expression in the sequenced tumor cells. **C,** The AP-1 network was inhibited with a dominant-negative Jun (JunDN) construct and a soft agar assay was used to determine colony forming potential. Representative images after crystal violet staining are shown. Scale bar = 5 mm. **D,** Quantification of the soft agar colony forming assay in (**D**). **E,** Percentage of live cells after JunDN transfection as determined by 7-AAD staining. **F,** UMAP plot of the untreated and chemotherapy-treated tumor samples showing distribution across the early and late cluster. **G,** Percent distribution of untreated and chemotherapy-treated samples by cluster (as in **F**). Significance determined by an unpaired student’s t-test where (*) P<0.05, (**) P<0.01, (***) P<0.001, (n.s.) P≥0.05.

### Genetic barcode lineage tracing system in human SCLC xenografts

To investigate tumor heterogeneity and chemoresistance in human SCLC cells, we again used genetic barcoding, but barcoded cells *in vitro* before injecting them as xenografts in immunocompromised mice. Like the *in situ* barcoding model (Fig. 1A), a barcoding retrovirus was designed that contains GFP, followed by the lineage barcode, and finally a polyA sequence (Fig. 3A). The inclusion of the polyA sequence at the 3’ end of the lineage barcode allows for best success at sequencing the barcodes using the 10X Genomics 3’ library prep. To be able to follow barcode populations over time, SCLC cell lines NCI-H209 (SCLC-A subtype) and NCI-H82 (SCLC-N subtype) were barcoded in culture and the cells grown as xenografts (Fig. 3B). A portion of the cells that were barcoded in culture were assessed by scRNA-seq (“pre-injection” sample), and a portion are used to generate xenografts. The initial xenografts that formed served as “pre-chemotherapy” samples, and upon dissection a portion are used for scRNA-seq, with the remainder serially transplanted into a new mouse. The mouse harboring this secondary xenograft was then treated with cisplatin and etoposide and tumors harvested and assessed by scRNA-seq to observe how chemotherapy treatment changes the transcriptional profile of these cells (“post-chemotherapy” sample). By this system, “pre-injection”, “pre-chemotherapy” and “post-chemotherapy” samples would contain matching barcodes giving insight into the clonal dynamics in these populations over time and treatment. Prior to the generation of xenografts, we sought to thoroughly profile the diversity of barcodes in the generated barcode retroviral pool. As this methodology starts with a split of the cells into the Pre-injection and xenograft samples, it is important to assess the doubling rates of these cell lines as clonal growth of singly labeled cells would be required for a significant overlap of the same barcodes between these samples. The doubling time of the two cell lines used to generate xenografts was therefore determined to be 2.02 days (NCI-H209) or 2.39 days (NCI-H82) (Fig. 3C). To determine the even distribution of unique barcoded cellular clones across two evenly divided samples, we used an Illumina targeted amplicon sequencing platform, to sequence the barcode region of the retroviral plasmid pool and the retroviral cDNA (Fig. 3D, 3E). To estimate the diversity of barcodes most accurately in the two samples, the PCR error rate in the non-variable region of the LBC was used to differentiate between unique barcodes and artifactual barcodes due to a PCR error (Fig. 3F). Finally, using the Chao2 asymptotic richness estimator (61, 62), to account for subsampling and PCR amplification errors (Supplementary Fig. S6), we determined the diversity of the barcode pool to be around 5,946 unique barcodes, well in excess of the 2,500 cells used to seed each xenograft (Fig. 3E). In order to ensure the barcodes captured in the pre-injection sample and the xenograft have sufficient overlap, prior to xenografting we profiled the barcodes in two independent samples taken from a population of barcoded cells at every doubling (Fig. 3C). As the number of cellular doublings increases, the fractional overlap of barcodes in two halves of a barcoded cellular population increases (48.3% overlap at 3 doublings), until a point at which the overlap is reduced (32.6% overlap at 5 doublings), presumably due to unequal growth of subclones or subclone death during culture (Fig. 3G). By assessing the barcode overlap over five doublings, the optimal number of doublings for sufficient overlap is identified to be three doublings (Fig. 3H).

**Figure 3.**
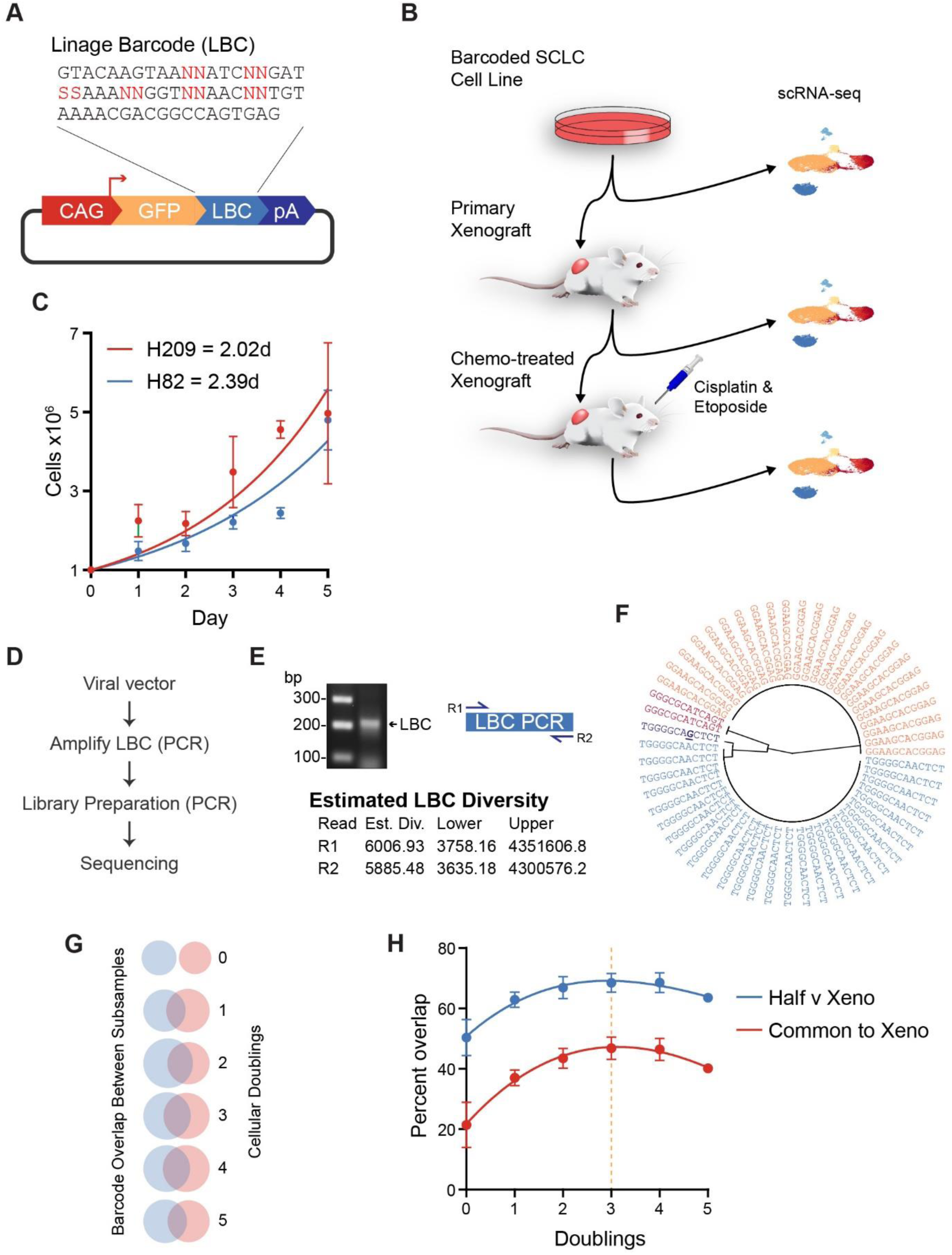
Generation and validation of barcoded xenografts and PCR error rate correction. **A,** The xenograft barcoding system. The barcode is inserted retrovirally, and contains a CAG promoter, GFP, the lineage barcode (LBC), and a polyA tail. **B,** Overview of the lineage tracing process. Cells are barcoded in culture and a portion taken for scRNA-seq. The remainder of the cells are injected as a xenograft. Half of this xenograft undergoes scRNA-seq, and the other half is injected as a serial xenograft into a new mouse, which is treated with chemotherapy and then undergoes scRNA-seq. **C,** Doubling time of the two SCLC cell lines used for making xenografts, NCI-H209 and NCI-H82 **D,** Process of sequencing LBCs for diversity validation. **E,** PCR purification of the LBC and subsequent sequencing. Diversity estimates from each read direction of the PCR products are shown. **F,** Radial plot showing the unbiased clustering of LBC similarity for a single sample. A single A>G transition is observed which is interpreted as a PCR error as its relative occurrence in the population is less than the calculated PCR error rate determined by variants in the LBC constant regions. **G,** Representation of the degree of overlap between two subdivided samples split at various cellular doublings measuring in (**C**). **H**, Degree of overlap between a single sample split in half to represent the labeled starting cell line and the cells to be divided into four subpopulations for xenograft injection.

### Transcriptomic profiling of tumor initiation and chemoresistance in SCLC xenografts

Xenografts were generated by injecting barcoded NCI-H209 or NCI-H82 cell lines into the hind flank of immunocompromised NOD-SCID mice, and growth was monitored over time (Supplementary Fig. S7, Supplementary Table S2). After dissection, tumors were sequenced via scRNA-seq and the remainder of cells were serially xenografted into a new mouse, which was treated with chemotherapy upon palpable xenografts were formed (36 days for NCI-H209 and 19.5 days for NCI-H82) and the subsequent chemotherapy-treated tumors were sequenced via scRNA-seq (Fig. 3A). While the bulk of these cells were of human origin, some cells from an immune or fibroblast mouse origin, as determined by mapping statistics between the human and mouse genome, were collected yet removed for subsequent analyses (Supplementary Fig. S8). The majority of the NCI-H82 (SCLC-N) xenograft cells, regardless of chemotherapy status cluster together, and there are very few changes in the global transcriptomes before and after chemotherapy (Fig. 4A, Supplementary Table S4). In contrast, the NCI-H209 (SCLC-A) post-chemotherapy cells display a robust transcriptomic shift from the pre-injection and pre-chemotherapy samplings, and cluster more closely with the SCLC-N tumors, indicating a shift to a more SCLC-N-like phenotype after chemotherapy (Fig. 4A, Supplementary Table S4). Both the tumor initiation and chemotherapy processes serve as cellular selection events, as the percentage of uniquely barcoded cells in each subsequent sample decreases nearly 10-fold at each stage, indicating only a subset of cells are able to initiate xenografts, and a smaller subset are able to survive chemotherapy and re-establish a tumor (Fig. 4B). However, similar to the *in situ* barcoding system, the overall percentage of sequenced LBCs driven by GFP expression was generally low (Supplementary Fig. S7). Therefore, we identified clonal populations in these cells by the identification of single nucleotide variants (SNVs) in the scRNA-seq data, focusing on the NCI-H209 cells as they showed greater dynamics upon chemotherapy exposure. Over time, the clonal distribution within the tumors shifts (Fig. 4C), indicating that this approach is amenable to identifying clonal subpopulations with phenotypic roles during SCLC tumor development. This xenograft study was designed with three unique serial transplantations from xenografts to post-chemotherapy xenografts which might each present with their own unique spontaneous SNVs, therefore we sought to further investigate the clonal dynamics over time in these three independent study arms (Fig. 4D, Supplementary Fig. S9, Supplementary Table S5). The three lineage arms show distinct modes of clonal evolution, but ultimately display increasing clonal diversity in the post-chemotherapy xenografts. This is reflected in the distribution of more disparate subclones in the chemotherapy-treated xenograft populations (Fig. 4E), yet all arising from the same subpopulations in the original cell line (Supplementary Fig. S9). To assess the relative plasticity of these subclonal populations, we divided the cells into multiple discreet clusters, and determined the degree of cluster residence at different stages of tumor development – i.e., from cell line to xenograft, or from xenograft to chemotherapy-treated xenograft (Fig. 4F). We observe that subclonal plasticity increases after exposure to chemotherapy (Fig. 4G).

**Figure 4.**
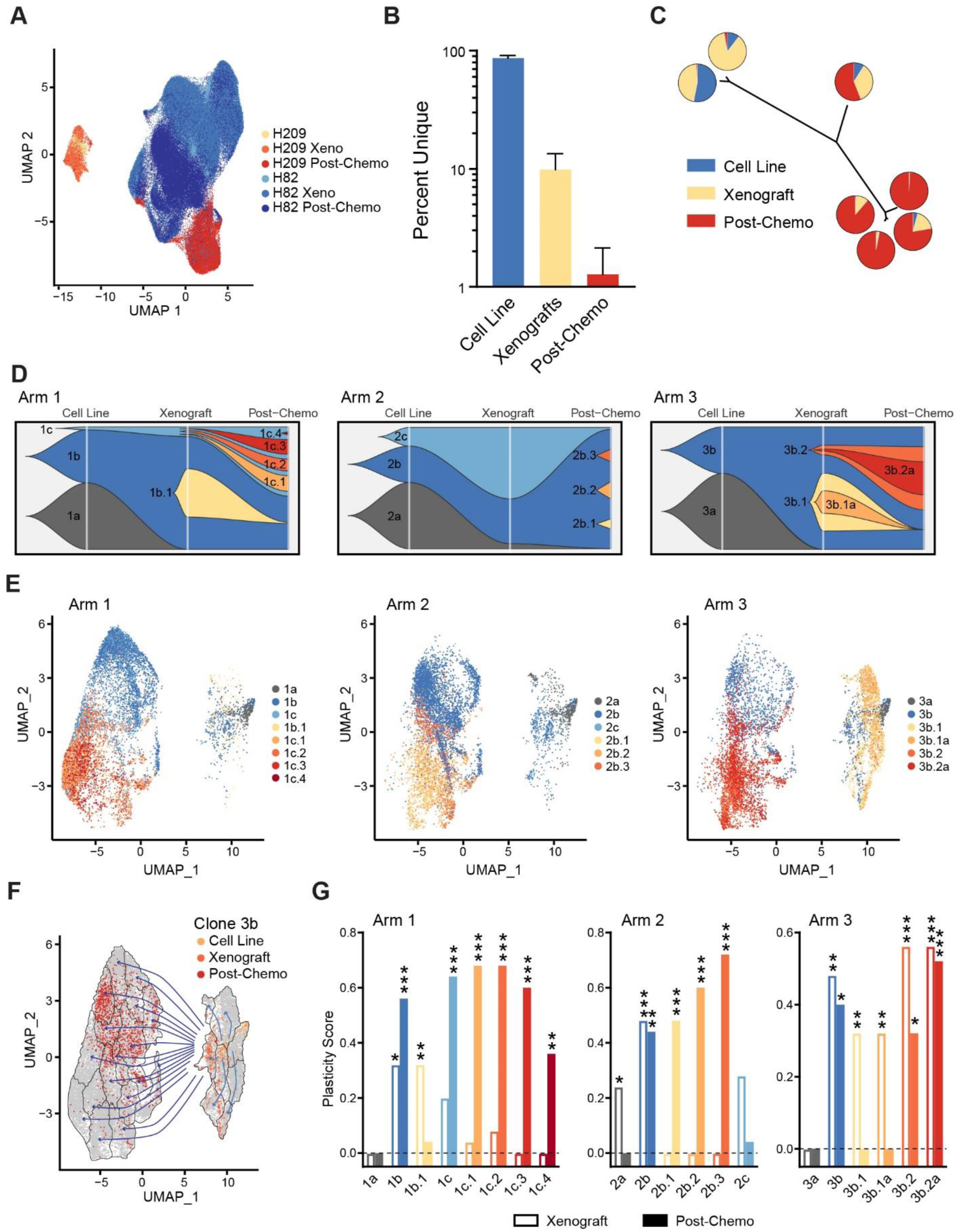
Transcriptomic plasticity and clonal evolution of SCLC xenografts. **A,** UMAP plot of all xenograft samples from the NCI-H209 and NCI-H82 samples. **B,** Percentage of unique LBCs sequenced at each stage of growth. Note log scale on y-axis. **C,** Individual subclones identified in the NCI-H209 cells showing a varying distribution amongst experimental stage. **D,** Muller plots showing subclonal evolution across the three independent serial transplants (arms) of this study, although the same starting cell line was used. **E,** UMAP showing the distribution of the subclones across the experimental stages, cell line (rightmost cells), chemo-naïve cells (Right cluster), and the chemo-treated cells (leftmost cluster). **F,** Schematic showing overclustering of the NCI-H209 data (dashed lines) and using Clone 3b as an example showing its positional changes and cluster residence between xenograft formation and chemotherapy treatment. **G,** Distance between the distribution of cells at either the xenograft or chemotherapy stages reported as a relative plasticity score. Scores and significance determined using a one-sided Kolmogorov– Smirnov test where (*) P<0.05, (**) P<0.01, (***) P<0.001, (n.s.) P≥0.05.

Further exploring the earliest subclones in original NCI-H209 cells, we clearly identify one subclone that does not contribute to xenograft formation (subclones 1a, 2a, and 3a), while the remaining subclones are found to populate the xenografts (Fig. 4D). We therefore identify these subclones (1b, 1c, 2b, 2c, and 3b) as tumor-initiating subclones (TICs) as it appears the cells capable of forming a tumor reside in these subclones. Differential gene expression between the cell line subclones readily distinguish TICs from non-TICs (Fig. 5A, Supplementary Fig. S10, Supplementary Table S6). As epithelial to mesenchymal transitions (EMT) is often associated with tumor invasiveness and growth we tested the relative EMT status of these clones and observe that EMT scores derived from the expression of known markers clearly separates TICs from non-TICs with the non-TICs possessing more of an epithelial state and the TICs displaying more of a mesenchymal state (Fig. 5B). This appears to closely relate to MYC activity as the non-TICs are enriched for genes associated with genetic amplification of *MYCL* and lack of amplification of *MYC* and *MYCN* (normalized enrichment score = 2.16, q-value = 2.4 x10^−4^), while the TICs show marked upregulation of genes associated with *MYC* and *MYCN* amplifications (normalized enrichment score = -3.47, q-value = <1 x10^−5^, Fig. 5C) (63). SCLC cells that are high in both EpCAM and CD24 and low for CD44 are enriched for long-term tumor propagating cells (64), and consistent with that discovery we observe that the TICs display similar characteristics in the starting cell line, however upon xenograft formation the levels of *EpCAM* are markedly reduced consistent with the acquisition of a more mesenchymal state (Fig. 5B, 5D). Furthermore, we observed in mouse SCLC tumors that the AP-1 complex is required for tumor development (Fig. 2), and in the TICs we also observe significant upregulation of JUN family members in growing xenografts, although there is a slight repression in the chemotherapy treated xenografts which differs from the mouse tumors (Fig. 5E).

**Figure 5.**
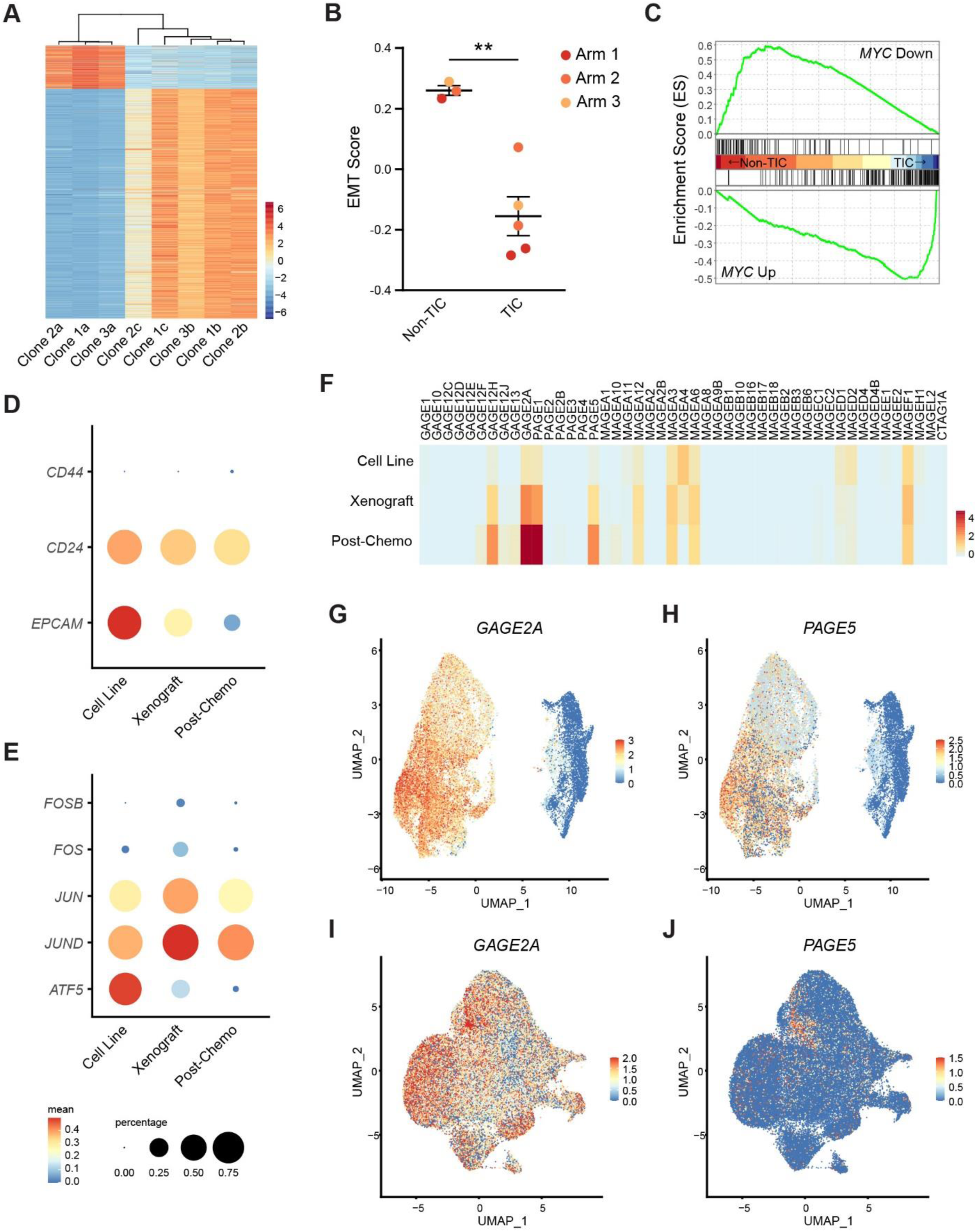
SCLC xenografts reveals tumor initiating clone characteristics and cancer testis antigens as markers of chemoresistance. **A,** Heatmap of differentially expressed genes found between the tumor initiating clones (TICs) and non-TICs. **B,** Relative EMT score of the TIC and non-TICs **C,** GSEA analysis comparing TIC and non-TICs identified strong enrichment of genes upregulated with *MYC* amplification in the TICs and genes downregulated by *MYC* amplification in non-TICs. **D,** TICs express markers of SCLC tumor propagating cells. **E,** TICs significantly express genes of the AP-1 complex, notable *JUN* and *JUND*. Scale bar and size reference is shared between **D** & **E**. **F,** Heatmap showing expression of the members of the CTA family. Expression of *GAGE2A* (**G** & **I**) and *PAGE5* (**H** & **J**) in both NCI-H209 cells (**G** & **H**) and NCI-H82 cells (**I** & **J**).

### Single-cell RNA sequencing reveals Cancer Testis Antigens as mediators of chemoresistance in SCLC

Intriguingly, the genes most upregulated upon chemotherapy treatment were the Cancer Testis Antigens (CTAs). CTAs are a large class of proteins almost exclusively expressed in male germ cells and many cancers where they act as oncogenes with key effects on proliferation, genomic stability, invasion, colony formation, and resistance to apoptosis (65–67). Of the broad class of CTAs members of the PAGE and GAGE families appear most upregulated upon chemotherapy treatment (Fig. 5F, Supplementary Table S7). We identified GAGE2A and PAGE5 (CT16) with the highest fold increase, however there is marked upregulation also of PAGE1 (Fig. 5E – 5J). We therefore sought to determine the role of CTAs in mediating response to chemotherapy. Biopsies from human SCLC patients, show immunopositivity for both GAGE2A and PAGE5, with more disperse staining of PAGE5 (Fig. 6A). In order to understand the requirement for CTA expression on chemoresistance, both *GAGE2A* and *PAGE5* were knocked down by retroviral expression of shRNAs in four SCLC cell lines (NCI-H209 and NCI-H1836, from the SCLC-A subtype; NCI-H82 and NJH29 from the SCLC-N subtype). While shRNAs were specifically designed to *GAGE2A* and *PAGE5*, due to the high degree of homology amongst CTA families, cross-reactivity is likely. Cells were treated with chemotherapy in culture (Supplementary Fig. S11) and the percentage of live cells was assessed via flow cytometry. SCLC-A type cell lines with both *GAGE2A* and *PAGE5* knocked down are nearly two-fold more sensitive to cell death (P = 0.013) caused by cisplatin (Fig. 6B), however knockdown in SCLC-N type cells showed a trend towards higher sensitivity yet this was not significant (P = 0.087) perhaps due to the inefficiency of the shRNAs to provide a large enough knockdown in the SCLC-N type cells which already possess high CTA expression (Fig. 5I, 5J). Subsequently, we tested if overexpression of *GAGE2A* or *PAGE5* via retroviral transduction altered response to cisplatin, etoposide, or combination cisplatin and etoposide. Cells with overexpression of either *GAGE2A* or *PAGE5* were significantly more resistant to cell death by chemotherapy in both SCLC-A cell lines (Fig. 6C) and SCLC-N cell lines (Fig. 6D) cells. Overexpression of *GAGE2A* or *PAGE5* had no impact on cellular survival in the absence of chemotherapy. To investigate the requirement for expression of *GAGE2A* and/or *PAGE5* for chemoresistance *in vivo,* xenografts were generated from the *shPAGE5*, *shGAGE2A*, or double knockdown SCLC cells, and the resulting xenografts treated with chemotherapy. Knockdown of *GAGE2A* or *PAGE5* had a modest impact on reducing growth of SCLC-A tumors not treated by chemotherapy (Fig. 6E). In SCLC-A xenografts treated with chemotherapy, knockdown of *GAGE2A*, *PAGE5*, or both had a robust increase in the duration of treatment response (Fig. 6F). In SCLC-N xenografts, which are generally more inherently chemoresistant (68), knockdown of *GAGE2A*, *PAGE5*, or both had an impact on growth in the absence of chemotherapy (Fig. 6G). Additionally, PAGE5 knockdown or the double GAGE2A and PAGE5 knockdown increased sensitivity to chemotherapy for the SCLC-N tumors, albeit to a lesser degree than the SCLC-A type cells due to incomplete knockdown of these CTAs (Fig. 6H). Overall, CTA knockdown significantly increased treatment response in human SCLC xenografts (Fig. 6F, 6G).

**Figure 6.**
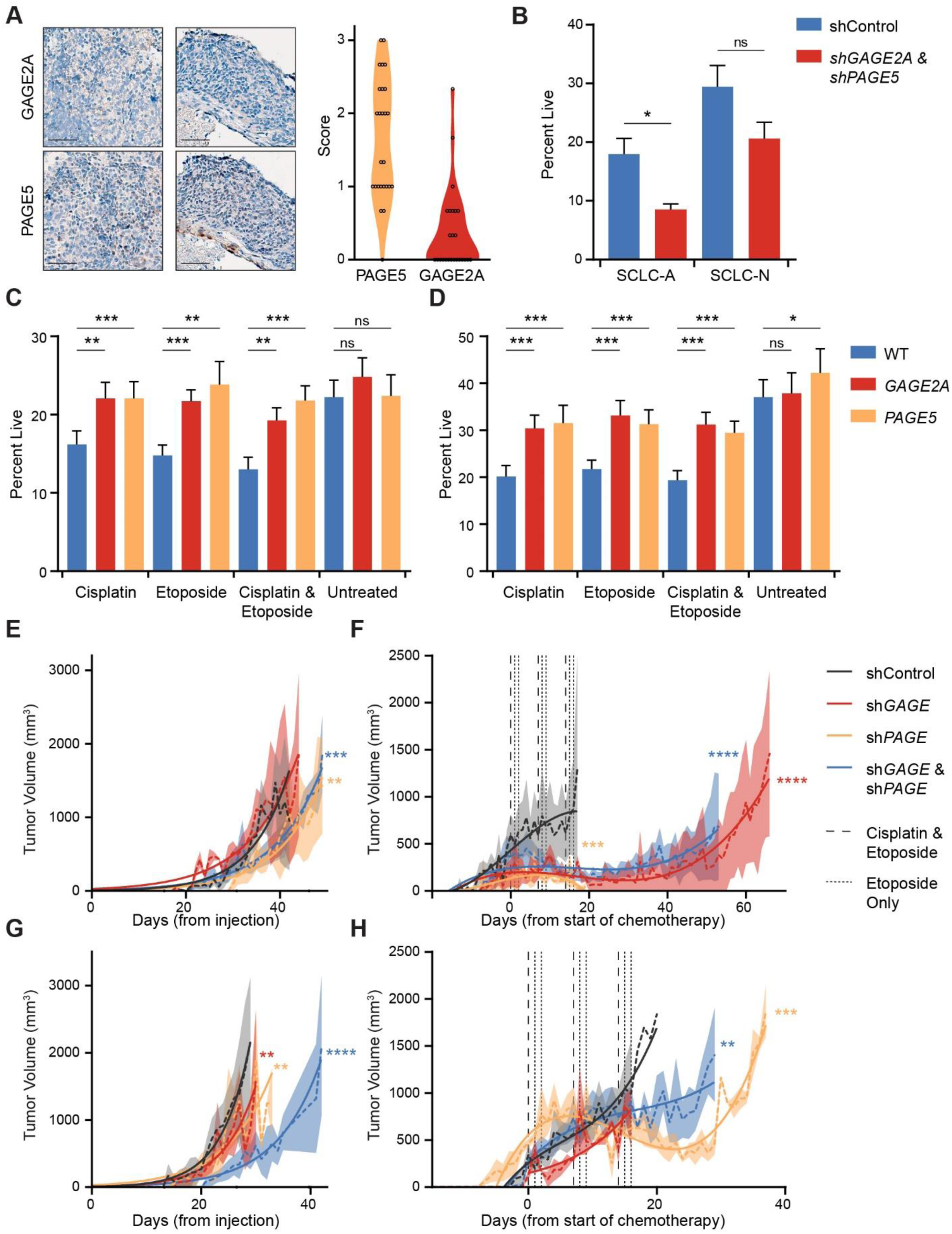
Cancer testis antigens are mediators of chemoresistance in SCLC. **A,** Staining of GAGE2A and PAGE5 in human SCLC patient samples. GAGE2A is surface expressed while PAGE5 is cytoplasmic. Scale bars = 50μm. Quantification on right shows the distribution score of each stain across the tumor sample. **B,** Quantification of living cells by Annexin V staining after treatment with cisplatin. Cell lines used were NCI-H1836 & NCI-H209 (SCLC-A) and NJH29 & NCI-H82 (SCLC-N). The percent living cells in SCLC-A cell lines (NCI-H1836 & NCI-H209) (**C**) and SCLC-N lines (NJH29 & NCI-H82) (**D**) after either cisplatin treatment, etoposide treatment, or treatment with both. Significance determined by an unpaired student’s t-test. Xenograft growth curves of SCLC-A (**E** & **F**) and SCLC-N (**G** & **H**) cell lines for untreated xenografts (**E** & **G**) and chemotherapy-treated xenografts (**F** & **H**). Significance determined by the extra sum-of-squares F test (*) P<0.05, (**) P<0.01, (***) P<0.001, (****) P≥0.0001.

## Discussion

SCLC is a devastating disease for which tumor heterogeneity has functional impacts for the treatment and course of the disease (21, 69, 70). Tumor heterogeneity over time and under chemotherapeutic pressure is not well understood, but single-cell technologies has emerged in recent years as a powerful tool for the understanding of tumor evolution. Here we use genetic barcode lineage tracing and subclone analysis to understand tumor dynamics in a mouse model of SCLC and human SCLC xenografts.

We performed scRNA-seq analysis on a cohort of RPR2 mice to investigate the genetic drivers of tumor development. We found the initial emergence of a population of neuroendocrine-high “early” tumor cells, which are maintained through the later stages of tumor development but make up proportionately less of the bulk of the tumor at later stages of tumor development. In their place, a “late” population of tumor cells arises and over later timepoints begins to be responsible for the majority of the tumor mass. Cells from chemoresistant tumors cluster most closely with this “late” tumor population. Members of the AP-1 network, especially *Jun* and *Fos* were highly upregulated in the late tumor cluster, and human SCLC cells in culture rely on the activation of the AP-1 network to maintain colony forming abilities. In SCLC xenografts, members of the AP-1 network are expressed in tumor initiating clones that also possess a more mesenchymal phenotype. JUN is capable of upregulating *Vimentin* and driving the epithelial to mesenchymal transition (71). Furthermore, this EMT transition is linked to metastasis (21, 72), which would be expected for a tumor initiating population of cells and a characteristic feature of human SCLC. Finally, there are many avenues for co-regulation of the AP-1 network and MYC either directly or mediated through the Jun kinase (JNK) (73–75).

Using a human xenograft model from two SCLC subtypes, SCLC-A and SCLC-N, we were able to investigate mechanisms of tumor development and treatment resistance. The SCLC-A xenografts show a profound transcriptomic shift after chemotherapy to a more SCLC-N-like profile consistent with high expression of *ASCL1* in chemotherapy-naive SCLC (68). In examining the barcode populations, two bottleneck events occur – the first in the generation of xenografts from cell lines, and the second in generating a chemoresistant xenograft, indicating that there are indeed discreet subpopulations of cells in an SCLC tumor capable of driving tumor initiation and chemoresistance. Subclonal analysis revealed that these two steps are not equally associated with cellular plasticity. While plasticity is increased slightly upon xenograft formation, the primary stage of enhanced plasticity is in response to chemotherapy (21). Furthermore, it is noteworthy that we observe both expansion of clones found in the original xenograft after chemotherapy treatment, and novel subclones that arise after chemotherapy. This indicates that while enhanced plasticity of tumor cells can lead to chemoresistance, many chemoresistant clones may reside in the tumor prior to treatment and then become unmasked upon the selective pressure of chemotherapy. Therefore, the mechanisms of evolution in SCLC tumors may not be uniform.

We, like Gardner *et al*. (76), have observed the upregulation of CTAs in chemo-treated SCLC. In particular, GAGE2A and PAGE5 are increased after chemoresistance in SCLC-A tumors and have relatively high expression in the inherently resistant SCLC-N tumors. To investigate the impact of GAGE2A or PAGE5 on mediating chemoresistance in SCLC, we knocked down *GAGE2A* and *PAGE5* in SCLC-A and SCLC-N cells and found increased sensitivity to cisplatin in culture. Knockdown of either GAGE2A, PAGE5, or both GAGE2A and PAGE5 led to increased sensitivity *in vivo* in SCLC-A and SCLC-N xenografts. Conversely, overexpression of GAGE2A or PAGE5 in SCLC cell lines led to increased resistance to chemotherapy *in vitro*. Finally, human SCLC biopsies show expression of GAGE2A and PAGE5 via immunostaining. PAGE5 has been identified to be expressed in some cancers, and in melanoma was elevated as an anti-apoptotic gene in response to platinum-based chemotherapy, where it was shown to be promote cell survival pathways (77). GAGE2A is another anti-apoptotic CT antigen, that seems to be related to treatment resistance in medulloblastoma (78). Another CTA, *CAGE*, has been implicated in the regulation of upregulation of Cyclin D and Cyclin E, resulting in G1 to S progression mediated by AP-1 (79). This cell cycle regulation was mostly mediated by *JUN* and *JUND*, in line with our observation that these are the AP-1 members most highly expressed in SCLC TIC subpopulations. CTAs have shown promise as potentially targetable, unique cancer antigens. For this reason, they make excellent candidates for immunotherapy such as CAR-T therapy and cancer vaccines (65-67, 80-83). Taken together, this indicates that the CTAs GAGE2A and PAGE5 are mediators of chemoresistance in SCLC and are expressed in human SCLC.

Initially we set out to perform single-cell DNA barcoding to trace ITH during SCLC development in xenografts and *in situ* using a mouse model. While our cellular labeling was robust in the xenograft model, it was much reduced in the Cas9-mediated *in situ* model due to the low efficiency of CRISPR recombination. Even still, our LBC detection levels were at best ∼12% in the xenograft model and undetectable in the *in situ* model. We surmise that this is a result of the relatively low expression on GFP in the transduced cells or the *in situ* model which is driven by the *Rosa26* locus expression. Future investigations may benefit from utilizing either a stronger promoter, or an inducible promoter to drive LBC expression. Furthermore, sequence capture techniques, such as the Feature Barcoding technology from 10X Genomics could be useful to enrich for LBC sequences. We did, however, demonstrate the PCR error is a considerable issue when performing genetic lineage barcoding as error introduced by PCR processing can artificially inflate the number of identified LBCs, and therefore confound downstream analyses of cellular heterogeneity by potentially subdividing unique clones. We therefore describe a framework where PCR error determined by the constant regions of an LBC can serve as threshold for any barcodes that appear at a lower frequency.

SCLC is a difficult disease to investigate with meager progress towards generating truly targeted therapeutics. The expansion of the single-cell clonal analysis has allowed us to examine ITH in SCLC with unprecedented resolution. Identifying the AP-1 network as being involved for SCLC tumor dynamics has provided knowledge of the critical early days of tumor formation in SCLC. Two potentially targetable antigens, GAGE2A and PAGE5 have been identified as direct mediators of chemoresistance in SCLC for the first time and may represent a new avenue for overcoming therapy resistance in SCLC in the future.

## Methods

### Ethics statement

Mice were maintained according to the guidelines set forth by the NIH and were housed in the Sanford Research Animal Research Center, accredited by AAALAC using protocols reviewed and approved by our local IACUC. The staining and scoring, and well as storage of the data for all human specimens was approved by the Sanford Health Institutional Review Board.

### Cell culture

SCLC lines NJH29 (H29), NCI-H82 (H82), NCI-H1836 (H1836), and NCI-H209 (H209) we used in this study. The cells were maintained in suspension and cultured in RPMI with 10% bovine growth serum and penicillin/streptomycin. All cell lines regularly tested negative for mycoplasma contamination and validated by IDEXX BioAnalytics. The NHJ29 cell line was a kind gift from Julien Sage.

### Generation and validation of barcoding libraries

#### Cloning of the barcoding libraries

To clone the retroviral barcoding library, a CAG-GFP retroviral plasmid was used as a backbone (gift from Fred Gage, Addgene plasmid #16664) (84). A poly-A sequence was subcloned from TetO-FUW-sox2, a gift from Rudolf Jaenisch (Addgene plasmid #20326) (85), to the 3’ end of the GFP at the PmeI site using InFusion Cloning (TaKaRa) and screened with PCR and restriction digests (Supplementary Table S8). DNA oligos containing the barcode sequence were ordered from Eurofins and annealed by heating to 95°C for five minutes and allowed to cool to room temperature over the course of several hours. The CMV-GFP-polyA plasmid was digested at PmeI and HindIII (added in the polyA cloning step), and the annealed barcode was inserted by InFusion cloning (TaKaRa) 3’ of the GFP and 5’ of the polyA sequence. Forty of the initial colonies were screened for presence of the barcode via PCR, and all four of the four sequenced were validated to contain the barcode using Sanger sequencing. Once presence of the barcode was confirmed in a number of colonies, the cloning product was transformed and plated to grow on five 15-cm plates of LB agar. Following overnight growth, all colonies were collected and pooled by flushing the plates with pre-warmed LB broth. Plasmid libraries were purified, and the product was pooled. Gamma-Retrovirus was produced by co-transfection of 293Ts with the barcoding retrovirus and retroviral packaging plasmid pCL-Amph. Retroviral supernatant was collected 48 and 72 hours after transfection and concentrated using the TaKaRa Retro-X concentrator.

To clone the AAV-LBC plasmid, a sgRNA targeted to *Rosa26* was subcloned from pU6-sgRosa26-1_CBh-Cas9-T2A-BFP, a gift from Ralf Kuehn (Addgene plasmid #64216) (86, 87) and inserted in to the AAV-KPL backbone (AAV:ITR-U6-sgRNA(Kras)-U6-sgRNA(p53)-U6-sgRNA(Lkb1)-pEFS-Rluc-2A-Cre-shortPA-KrasG12D_HDRdonor-ITR (AAV-KPL), a gift from Feng Zhang (Addgene plasmid #60224) (88) at SacI and MulI using InFusion cloning (TaKaRa), and screened using PCR and Sanger sequencing. Left and right homology arms to *Rosa26,* as well as a polyA signal were subcloned from pR26 CAG/GFP Dest, a gift from Ralf Kuehn (Addgene plasmid #74281) (86, 87) in to the AAV9-r26 plasmid at PmlI. The barcode library, attached to a GFP was ordered from ThermoFisher’s GeneArt program, and was inserted in to the AAV9-r26 plasmid at PmlI and BamHI via InFusion cloning. After the first 30 colonies were screened for insertion of the barcode by restriction digest, PCR, and Sanger sequencing, chemically competent cells were transformed with the plasmid pool and plated on ten 10-cm LB plates. After overnight growth, all plates were washed with pre-warmed LB and the pooled colonies were purified. The AAV-LBC plasmid was used to generate AAV9 at the University of Michigan Viral Vector Core.

#### Validation of barcode diversity

To profile the diversity in the barcode pools, targeted amplicon sequencing was performed on the barcode region. PCR was used to amplify the barcode and add partial adaptors for Illumina sequencing. The minimal number of cycles needed to amplify the barcode region was used to minimize the risk of introducing variants or over-saturate the sample with a limited number of barcodes that had been disproportionately amplified. Samples were sequenced on an Illumina platform at Genewiz. The targeted amplicon sequencing of the barcodes was trimmed, and QC performed via CutAdapt. GREP (bash) was used to identify barcodes and export them to R Studio for analysis. The true number of barcodes was determined by finding the optimal number of clusters in a similarity based hierarchical clustering model that minimizes the distance in the PCR error rates between the random regions of the barcode and the constant region of the barcode. The total number of unique barcodes was estimated using Chao2 asymptotic richness estimator in R (61).

#### Doubling time assays to determine the fractional overlap of barcodes

In order to ensure sufficient overlap in barcodes between the “pre-growth” sample and xenografts, we sought to determine the optimal number of doublings before sufficient overlap in two independent samples was observed. 1.6M cells were seeded and barcoded using the CAG-GFP-BC retrovirus and a spinfection at 940 xg for 2 hours. At each doubling for five doublings following spinfection, one well of cells was harvested and split in two. The barcode region was amplified off the cDNA, and partial adapters for Illumina-based sequencing was added. Amplicon sequencing was performed at Genewiz (South Plainfield, NJ) using an Illumina miSeq platform. The barcodes were analyzed using a custom R script after trimming and QC via CutAdapt. To verify the results of the barcode sequencing, computational modeling was used. We simulated the same doubling time experiment 1,000 times using a custom R package. Based on the results of the sequencing, modeling, and previously published work (36), three doublings after barcoding gives sufficient overlap between two independent samples and will be used to generate the barcoded xenografts.

### Mouse protocols

#### In vivo model

The well-characterized *Rb^lox/lox^, p53^lox/lox^, p130^lox/lox^* SCLC mouse model (46) was bred to the *H11^lox-stop-lox-Cas9^* mouse (47) model (Jax #026816) to generate the RPR-Cas9 mouse used in this work. Tumors were initiated by intratracheal injection with Ad-CMV-Cre (Baylor Viral Vector Core) to delete the *Rb, p53,* and *p130* loci, and induce expression of Cas9. At one-month intervals for five months after tumor initiation, the AAV-LBC virus was delivered to separate cohorts of mice via intratracheal instillation (89) to barcode the forming tumors at the *Rosa26* locus using the CRISPR-Cas9 system. Mice were euthanized at one-month intervals after their Ad-CMV-Cre injection, up to six months, or when they became moribund according to institutional IACUC guidelines. Two mice received chemotherapy at 5 mg/kg cisplatin and 10 mg/kg etoposide on day one, and 10 mg/kg etoposide on days two and three with subcutaneous saline, repeated for three weeks, and were allowed to progress until they were moribund (24).

#### Xenografting

NCI-H209 and NCI-H82 SCLC cell lines were validated as pathogen free (IDEXX BioAnalytics) and then barcoded with the rCMV-GFP-BC retrovirus by spinfection at 940 xg for 2 hours. Each barcoded line was allowed to double three times, to ensure overlap in barcodes in the cells sampled and cells used for xenografting. Immediately prior to xenografting, a portion of the cells were removed to generate the single-cell RNA sequencing library (“pre-growth” sample). To make the xenografts, 2500 cells were mixed in a 1:1 ratio with matrigel (Corning Life Sciences) and injected into the hind flank of NOD-SCID mice. Xenografts were measured daily after the tumors were palpable by hand in the hind flank. After xenografts reached 3 cm^3^ total volume, mice were euthanized, and the xenografts harvested. Tumors were dissected, dissociated, and a portion of the cells were used for single-cell RNA sequencing (“pre-chemotherapy” sample). The remainder of cells were mixed in a 1:1 ratio with matrigel and injected into a new NOD-SCID mouse. The mice that received these serial xenografts received chemotherapy at 5 mg/kg cisplatin and 10 mg/kg etoposide on day one, and 10 mg/kg etoposide on days two and three, repeated for three weeks after tumors were palpable to generate chemoresistant xenografts (24). When these mice reached a total tumor burden of 3 cm^3^, they were euthanized, and the tumors were dissociated and used to generate single-cell RNA sequencing libraries (“post-chemotherapy” sample). Xenograft growth was plotted by determining the best fit between an exponential growth model or a cubic polynomial model by Akaike’s Information Criteria (AICc). Significance was determined by the extra sum-of-squares F test.

### Tumor profiling using single cell RNA sequencing

#### Tissue processing and library preparation

The “pre-growth” from the xenograft model were prepared for scRNA-seq according to the 10X Genomics protocols. After xenografts reached a cumulative volume of 2 cm^3^, mice were euthanized, and tumors dissected. The MACS (Miltenyi Biotec) human tumor dissociation kit was used to digest the tumors, and cells were prepared for scRNA-seq using the 10X Genomics protocols. After the RPR-Cas9 reached their endpoint, they were euthanized via cervical dislocation and lungs were harvested. Tissues were prepared for flow cytometry according to the 10X Genomics protocols, using the MACS mouse tumor dissociation kit. After dissection, the lungs were sorted for GFP+ cells, which are the barcoded population, and cells were prepared for scRNA-seq according to the 10X Genomics protocol. Tumors were micro-dissected from three mice by cutting out one tumor lesion, which was sequenced without undergoing FACS to capture the stromal and microenvironment cells. A 10X Genomics Chromium Controller was used for the library preparation of all tumors.

#### Tumor Histology

One lobe of the lung and one lobe of the liver from each of the RPR-Cas9 mice were taken for histology to verify the presence of tumors in this sample. Samples were fixed in 4% paraformaldehyde for 15 minutes and transferred to 30% sucrose for 24-48 hours. The samples were then embedded and cryosectioned before staining. Tumors were stained for GFP and Cas9 to confirm that the barcoding system was successfully induced by the Adenovirus induction of Cas9 expression and AAV9 induction of GFP expression.

### Informatics approach

The scRNA-seq data was processed uing 10X Genomics CellRanger software (10X Genomics, version 6.0.1). Initially CellRanger count was run on each sample followed by CellRanger aggregate to generate one expression matrix, a vector of genes and a vector of barcodes. CellRanger was run with the *–expect-cells* parameter set to the expected number of cells. The expression matrix and the vectors of genes and barcodes were imported to R and analyzed using the package Seurat (90) (version 4.1.1) to find cell clusters and their markers. Prior to clustering cells were further filtered based on percent mitochondrial RNA and number of genes expressed. Monocle3 (90–93) (version 1.2.9). was used similarly to both cluster cells and to run psuedotime analysis. Unbiased cell type determination was performed using Garnett (94) (version 0.2.8) using the provided mmLung_markers.txt classifier.

For the analysis of clonal evolution of cells and retrieval of LBC from cells, sample level scRNA fastq data was partitioned into per-cell fastq. First cell barcodes from cells that were deemed to be viable in the CellRanger were retrieved using Seurat. These cell barcode were used to simplify the demultiplexing step and to assign reads to cells using cutadapt (95) (version 2.0). Each cell specific fastq was mapped to human and to mouse genome using STAR (96) (version 2.7.10a) and the percentage mapping rate was used to identify and filter mouse cells. Presence of LBC in the reads of each cell was checked by using cutadapt to capture reads that possess the constant flanking regions of the LBC. Per cell SNPs and Indels were called using GATK best practices (97). To speed up the analysis and avoid computational limitations, haplotypeCaller was run in the GVCF mode followed by GenomeicDBImport and GenotypeGVCFs for each gene that was expressed in the scRNA data. Per gene vcf files were finally merged using bcftools (98) (version 1.9) to produce a joint vcf for all cells and genes. Cell relatedness and clonal evolution was investigated using Dendro package in R (99). Muller plots were designed using Fishplot (100) (version 0.5). Gene set enrichment analysis was performed using GSEA (101) (version 4.2.3). EMT scores defined by imogimap (102) (version 0.0.0.9000)

### Validation of candidates

#### Chemotherapy treatment of cells in culture

An alamar blue assay was used to determine the IC_50_ value of cisplatin and etoposide in NCI-H82 and NCI-H209 cell lines. Cells were seeded in 96 well plates and treated with either drug, with concentrations spanning three orders of magnitude. Cellular viability was assessed daily via Alamar Blue. The IC_50_ for Cisplatin was determined to be 2.876 μM. The IC_50_ for etoposide was determined to be 0.110 μM. These values were used for the resulting experiments. SCLC-N lines H29 and H82, and SCLC-A lines NCI-H1836 and NCI-H209 were treated with cisplatin and etoposide at the IC_50_ values for three days at cycles resembling the in vivo chemotherapy treatment (Cisplatin and etoposide day 1, etoposide only days 2 and 3). Cells were harvested ono days two and three and RNA was extracted following the Trizol (Ambion Biosciences) protocol, and RNA was converted to cDNA using the NEB ProtoScript Reverse Transcriptase Kit. Expression levels of *PAGE5* and *GAGE2A* were quantified via qPCR.

#### Knockdown of GAGE2A and PAGE5

shRNAs targeting *PAGE5* and *GAGE2A* (Supplementary Table S3) were designed using pSicoligoMaker3 (Ventura lab). shRNA oligos were cloned in to the lentiviral backbone pSicoR, a gift from Tyler Jacks (Addgene plasmid #11579) (103). The pSicoR-shPAGE5 and pSicoR-shGAGE2A were made into a second-generation lentivirus using pMD2.G and psPAX2 as packaging plasmids and concentrated overnight using the TaKaRa Retro-X retroviral concentrator. SCLC-N lines H29 and H82, and SCLC-A lines H209 and H1836 were infected with pSicoR-shPAGE5 or pSicoR-shGAGE2A and sorted by GFP expression using the BD FACS Jazz. Knockdown of *PAGE5* and *GAGE2A* expression was validated in the sorted cells with qPCR. To generate a double-knockdown, the sorted cells were infected with the reciprocal virus and expression of both *PAGE5* and *GAGE2A* was assessed via qPCR. The single- and double-knockdown cells were used to generate xenografts to investigate the dependence of the chemoresistance phenotype on *PAGE5* or *GAGE2A* expression. 150,000 cells were mixed in a 1:1 ratio with GelTrex (Gibco) and injected in to the hindflank of NOD-SCID mice. Due to supply chain disruptions, a switch from Matrigel to GelTrex was necessary, however they both function the same way in providing some extracellular matrix to aid in xenograft injection. Tumors were allowed to grow until a cumulative volume of 3 mm^3^ was reached, at which point mice were euthanized and the tumors kept for histology to validate the knockdown of *PAGE5* and *GAGE2A*. Half of the mice were treated with chemotherapy at 5 mg/kg cisplatin and 10 mg/kg etoposide on day one, and 10 mg/kg etoposide on days two and three, repeated for three weeks after tumors were palpable. In culture, the *shPAGE5*, *shGAGE2A*, and double knockdown cells were treated with the IC_50_ value of cisplatin and viability was assessed via Annexin V and propidium iodide staining by flow cytometry (Biolegend APC Annexin V Apoptosis Detection Kit).

#### Overexpression of PAGE5 and GAGE2A

*GAGE2A* and *PAGE5* overexpression retroviruses were generated by amplifying the transgenes from SCLC cell lines and cloning them into the CAG-GFP retroviral backbone. NJH29, NCI-H82, NCI-H1836, and NCI-H209 cells were transfected with either the rCAG-PAGE5-GFP or rCAG-GAGE2A-GFP vectors by spinfection with concentrated virus at 940xg for two hours. Transduced cells were treated with the IC_50_ value of cisplatin, etoposide, or cisplatin and etoposide assessed for response to chemotherapy by Annexin V and propodium iodide staining, and efficiency of overexpression assessed by qPCR.

#### Knockdown of the AP-1 pathway via overexpression of dominant-negative Jun

To inhibit the AP-1 complex, cJun was knocked down by transfection with a dominant-negative cJun construct. This is a common method for inhibiting the formation of the Jun/Fos AP-1 complex (53, 60). pMIEG3-JunDN was a kind gift from Alexander Dent (Addgene plasmid #40350) (60). NCI-H82, NJH29, NCI-H1836, and NCI-H209 SCLC cell lines were transfected with pMIEG3-JunDN using Lipofectamine 3000. Upon visual GFP detection, cells were sorted using FACS for GFP expressing cells. Cells were seeded in to 6 well plates for a soft agar colony formation assay. Briefly, 0.8% Seaplaque agar (Lonza) was used as a bottom layer and 10,000 cells per well were seeded in 1.2% agar in the top layer. Plates were fed with full RPMI as needed to prevent drying out. 10 days after seeding, colonies were observed by eye and the plates were stained with 0.001% crystal violet for one hour, and plates were photographed. The number of crystal violet colonies stained was quantified with a custom CellProfiler script.

### Analysis of PAGE5 and GAGE2A expression in human SCLC biopsies

#### Staining of human biopsies

Human SCLC biopsies were obtained from the Sanford Health Biobank. Slides were stained with anti-PAGE5 (Invitrogen PA5-50470) or anti-GAGE2A (Aviva Systems ARP64957-P050). Stained slides were scanned using an Apereo AT2 slide scanner. Three independent researchers viewed and scored the scanned slides based on positivity, distribution, and intensity of staining. Positivity was a binary score, with the sample earning a positive score from any singular positive cell. Distribution was scored on a 0-3 scale: 0 for 0-5% of the sample staining positively, 1 was assigned to samples 6-30% positive, 2 for samples 31-60%, and 3 for samples more than 60% positive for PAGE5 or GAGE2A. For staining intensity, a score 0-3 was assigned. 0 for samples with no PAGE5 or GAGE2A staining, 1 for samples with light staining, 2 for samples with moderate staining, and 3 for samples with intense staining.

## Acknowledgements

We would like to acknowledge the NIH NIGMS Center for Cancer Research P20GM103548 for pilot award support and the NIH NCI/NIGMS grant R01CA233661 for research support (to M.S. Kareta). The NIH NCI grant 5F31CA243149 for training and research support (to H.Wolenzien). The Sanford Research Functional Genomics and Bioinformatics Core is supported by the NIGMS Center for Pediatric Research 5P20GM103620, and the Pathology and Flow Cytometry Cores are supported by the NIGMS Center for Cancer Research P20GM103548. H. Wollenzien is thankful to the University of South Dakota-Neuroscience, Nanotechnology and Networks (USD-N3) Program, supported by a grant from the National Science Foundation Research Traineeship program DGE-1633213.

## Author Contributions

**H. Wollenzien:** Conceptualization, resources, data curation, formal analysis, validation, investigation, visualization, methodology, writing–original draft, writing–review and editing

**Y. Afeworki:** Data curation, software, formal analysis, writing–review and editing

**R. Szczepaniak-Sloane:** Formal analysis, writing–review and editing

**A. Restaino:** Formal analysis, writing–review and editing

**M.S. Kareta:** Conceptualization, resources, data curation, software, formal analysis, supervision, funding acquisition, validation, investigation, visualization, methodology, writing– original draft, project administration, writing–review and editing.

---

**Supplementary Table S1.** Description of samples in the *in situ* study.

| <u>Sample ID</u> | <u>Harvest Time</u> | <u>Barcode Time</u> | <u>Chemotherapy</u> | <u>Notes</u> |
| --- | --- | --- | --- | --- |
| HW1603 | 2 mo. | 1 mo. | - | - |
| HW1604 | 3 mo. | 1 mo. | - | - |
| HW1602 | 4 mo. | 1 mo. | - | - |
| HW1656 | 4 mo. | 2 mo. | - | - |
| HW1614 | 4 mo. | 3 mo. | - | - |
| HW1641 | 5 mo. | 0 mo. | - | - |
| HW1553 | 5 mo. | 2 mo. | - | - |
| HW1640 | 5 mo. | 3 mo. | - | - |
| HW1672 | 6 mo. | 1 mo. | cis/etop | - |
| HW1673 | 6 mo. | 2 mo. | - | harvest 5.5 mo. |
| HW1645 | 6 mo. | 3 mo. | - | harvest 5.5 mo. |
| HW1534 | 7 mo. | 2 mo. | cis/etop | - |

**Supplementary Table S2.** Description of samples in the xenograft study.

| <u>Sample ID</u> | <u>Parental Cell Line</u> | <u>Cell Growth</u> | <u>Chemotherapy</u> | <u>Arm</u> | <u>Notes</u> |
| --- | --- | --- | --- | --- | --- |
| HW_H209 | NCI-H209 | Cell Line | - | 1,2,3 | - |
| HW_L1419 | NCI-H209 | Xenograft | - | 1 | - |
| HW_L1420 | NCI-H209 | Xenograft | - | 3 | - |
| HW_R1419 | NCI-H209 | Xenograft | - | 2 | - |
| HW_R1420 | NCI-H209 | Xenograft | - | - | No 2° transplant |
| HW_T1620 | NCI-H209 | Xenograft | cis/etop | 1 | - |
| HW_T1623 | NCI-H209 | Xenograft | cis/etop | 2 | - |
| HW_T1624 | NCI-H209 | Xenograft | cis/etop | 3 | - |
| HW_H82 | NCI-H82 | Cell Line | - | 1,2,3,4 | - |
| HW_L1422 | NCI-H82 | Xenograft | - | 1 | - |
| HW_L1424 | NCI-H82 | Xenograft | - | 3 | - |
| HW_R1422 | NCI-H82 | Xenograft | - | 2 | - |
| HW_R1424 | NCI-H82 | Xenograft | - | 4 | - |
| HW_T1426 | NCI-H82 | Xenograft | cis/etop | 2 | - |
| HW_T1429 | NCI-H82 | Xenograft | cis/etop | 1 | - |
| HW_T1430 | NCI-H82 | Xenograft | cis/etop | 3 | - |
| HW_TL1421 | NCI-H82 | Xenograft | cis/etop | 4 | Left Xenograft |
| HW_TR1421 | NCI-H82 | Xenograft | cis/etop | 4 | Right Xenograft |

**Supplementary Figure S1.**
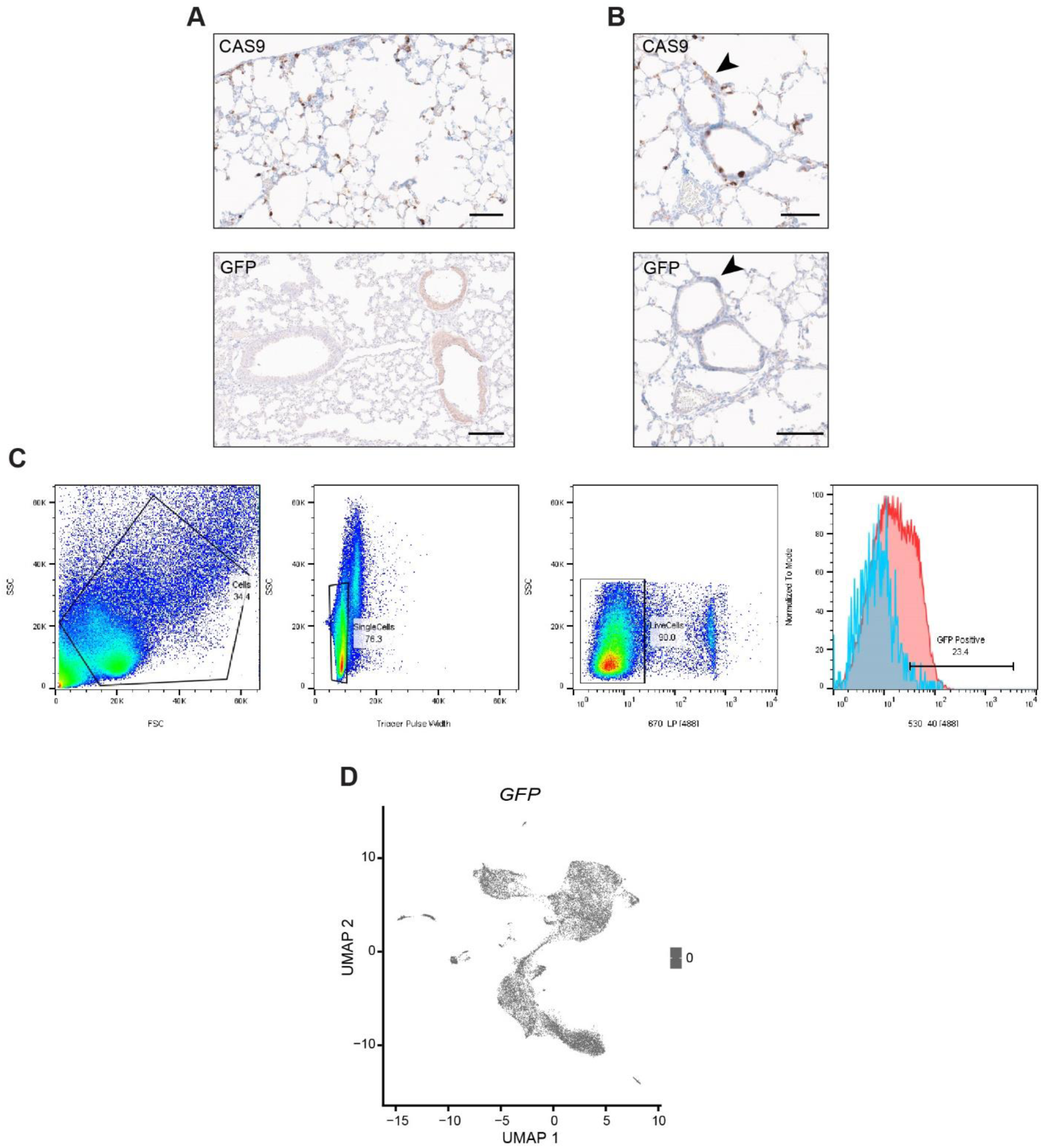
Validation of *in situ* barcoding model. **A,** Separate lung sections displaying positivity for both CAS9 and GFP. **B,** Co-staining of representative sections showing colocalization of both CAS9 and GFP. **C,** Flow-sorting strategy for isolating GFP+ cells from harvested lungs transduced with Adeno-Cre and the AAV-LBC viruses. **D,** UMAP plot displaying expression of GFP in the *in situ* scRNA-seq data.

**Supplementary Figure S2.**
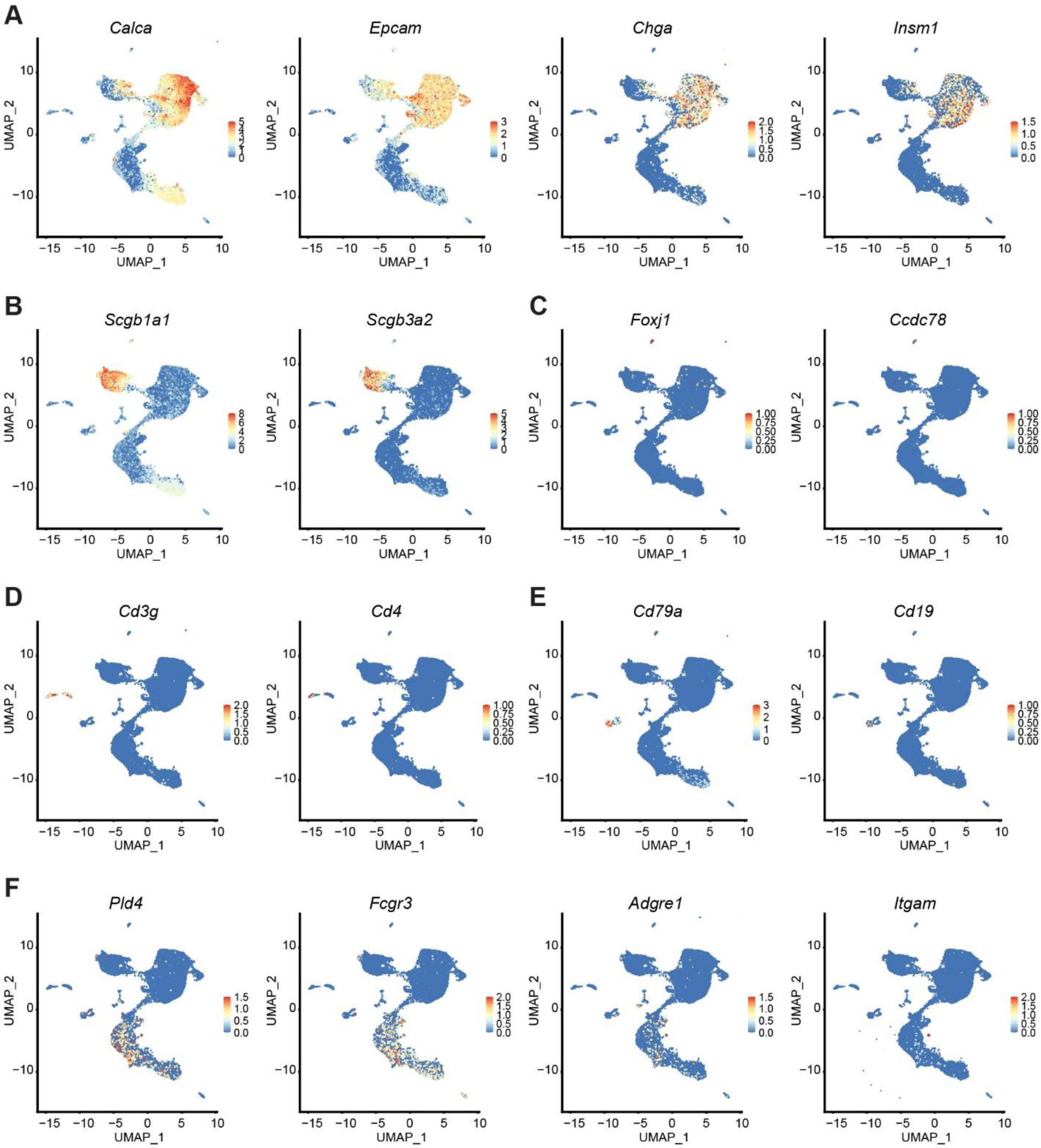
Characterization of cell types isolated from the *in situ* scRNA-seq data. Marker expression for SCLC (**A**), club cells (**B**) ciliated cells (**C**) T-cells (**D**) B-cells (**E**) myeloid cells (**F**).

**Supplementary Figure S3.**
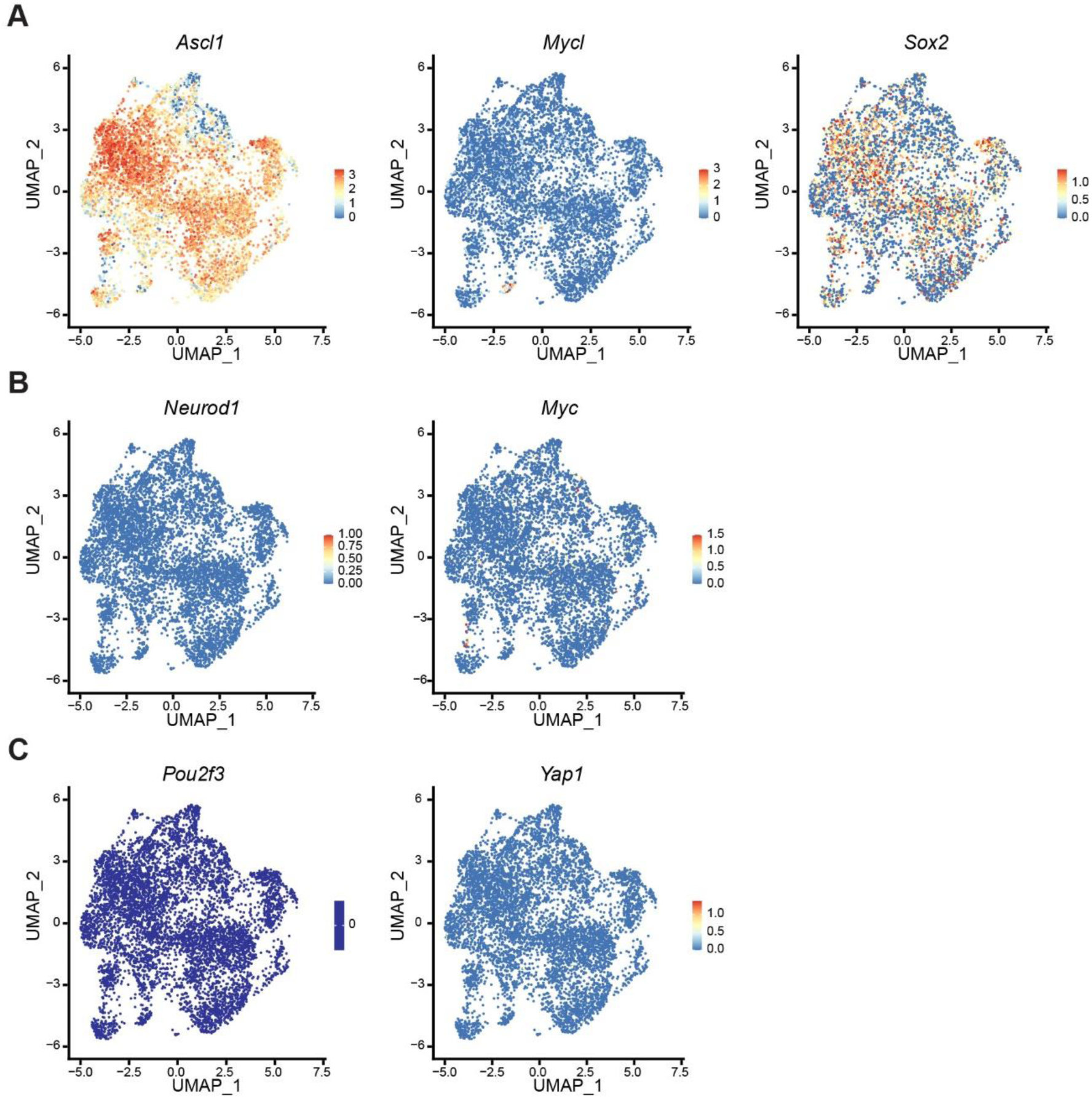
SCLC Subtype-specific marker expression from the RPR2-Cas9 mouse model. **A,** Markers for the SCLC-A subtype including *Ascl1*, *Mycl*, and *Sox2*. **B,** SCLC-N subtype markers including *Neurod1* and *Myc*. **C,** Expression of non-neuroendocrine SCLC markers *Pou2f3* and *Yap1*.

**Supplementary Figure S4.**
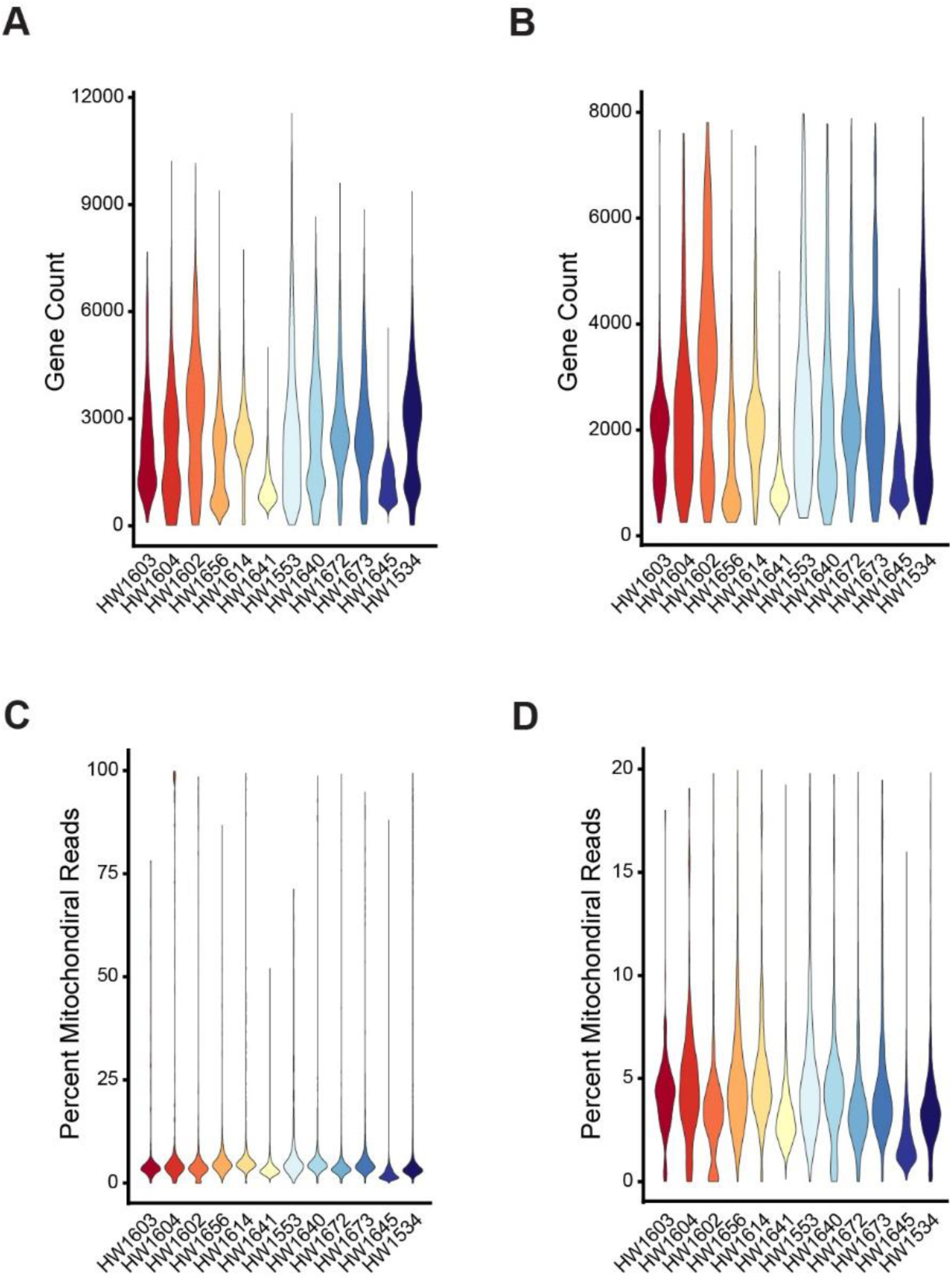
Gene counts and filtering. Gene counts for all sequenced cells (**A**) and after filtering (**B**) Percent mitochondrial counts per cell for all cells sequenced (**C**), and post-filtering (**D**).

**Supplementary Figure S5.**
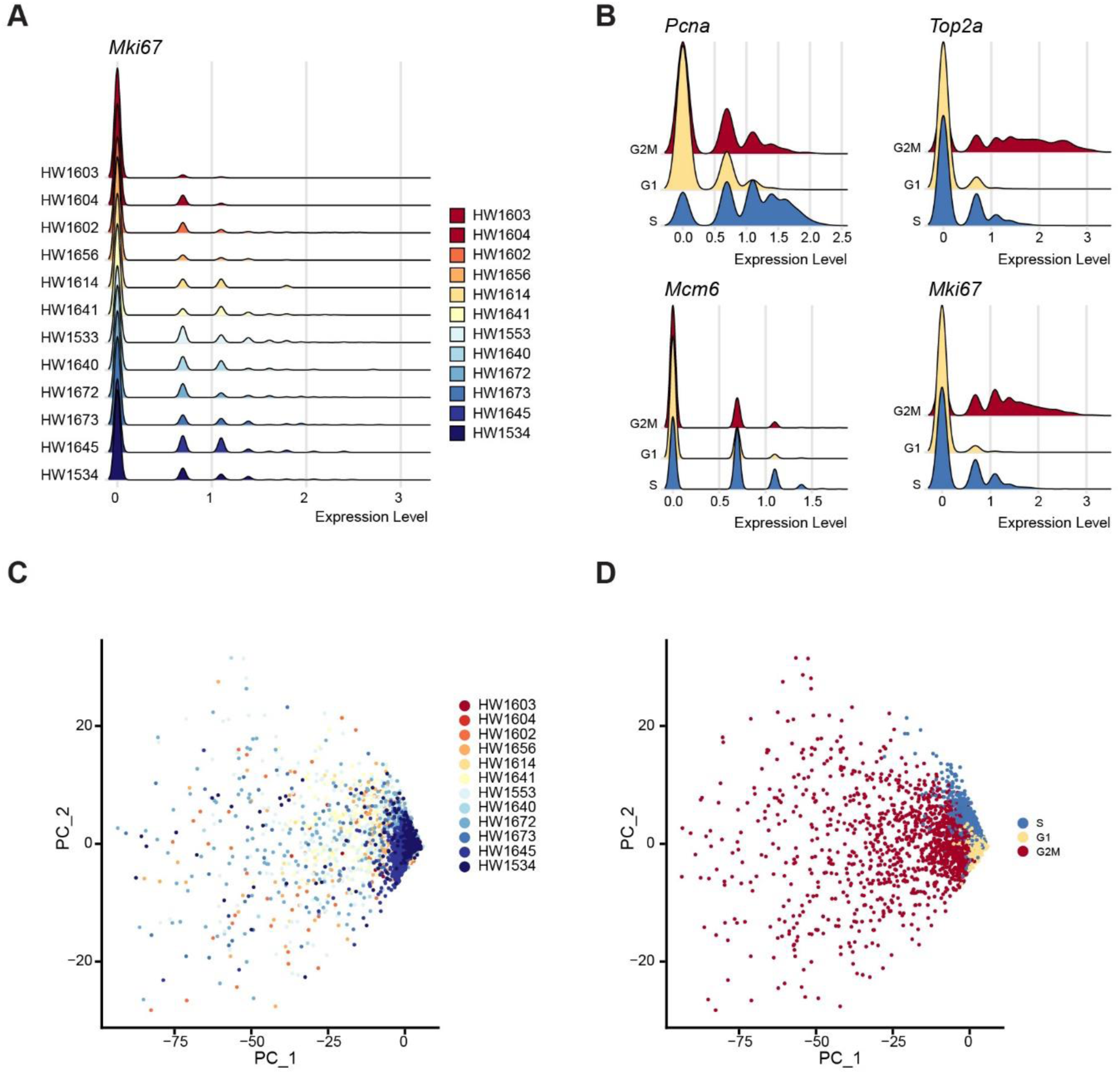
Cell cycle stage assignment of the SCLC cells from the *in situ* model. **A,** *Mki67* expression of the SCLC cells. **B,** Reliability of the cell cycle stage assignment by the markers (*Pcna*, *Top2a*, *Mcm6*, *Mki67*). Principal component plot of the *in situ* SCLC cells labeled by (**C**), and by cell cycle stage assignment (**D**).

**Supplementary Figure S6.**
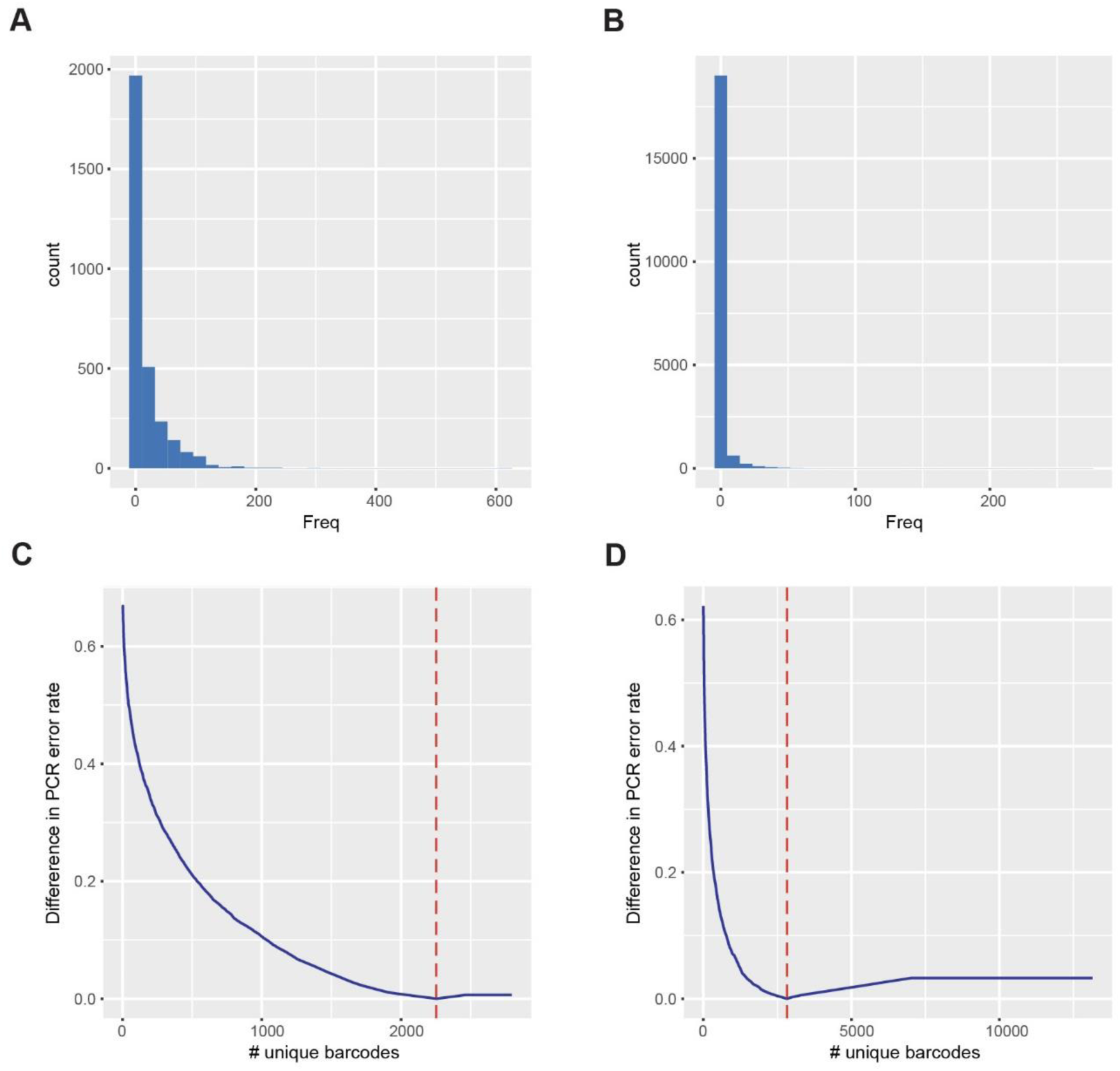
Determination of LBC counts by PCR error correction. Histogram of barcode frequency identified in the purified LBC vector (**A**) and the LBC vector packaged into retroviral particles (**B**). Estimation of minimal PCR error rate to determine corrected LBC count in the LBC vector (**C**) and the LBC vector packaged into retroviral particles (**D**).

**Supplementary Figure S7.**
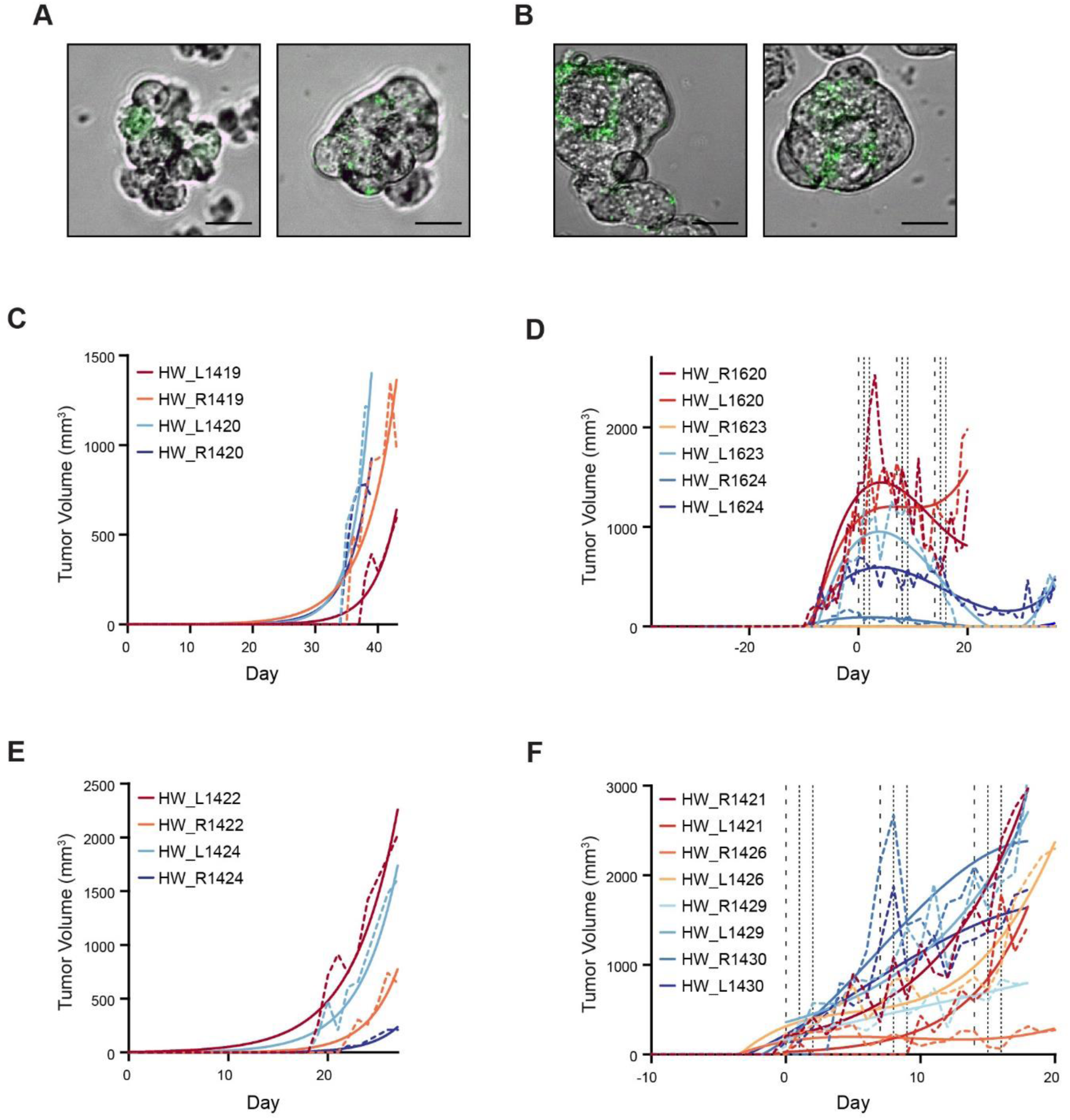
Validation of the LBC labeling and xenograft growth. Confocal imagery of the NCI-H209 (**A**) and NCI-H82 (**B**) cell lines expressing GFP after transduction with the Retro-LBC virus. Scale bars = 20 μm. Growth of the NCI-H209 xenografts in the primary xenograft without treatment (**C**) and the secondary xenograft during chemotherapy treatment (**D**). Growth of the NCI-H82 xenografts in the primary xenograft without treatment (**E**) and the secondary xenograft during chemotherapy treatment (**F**). Dashed lines represent the measured size of the xenografts and the solid line is the best-fit model of xenograft growth.

**Supplementary Figure S8.**
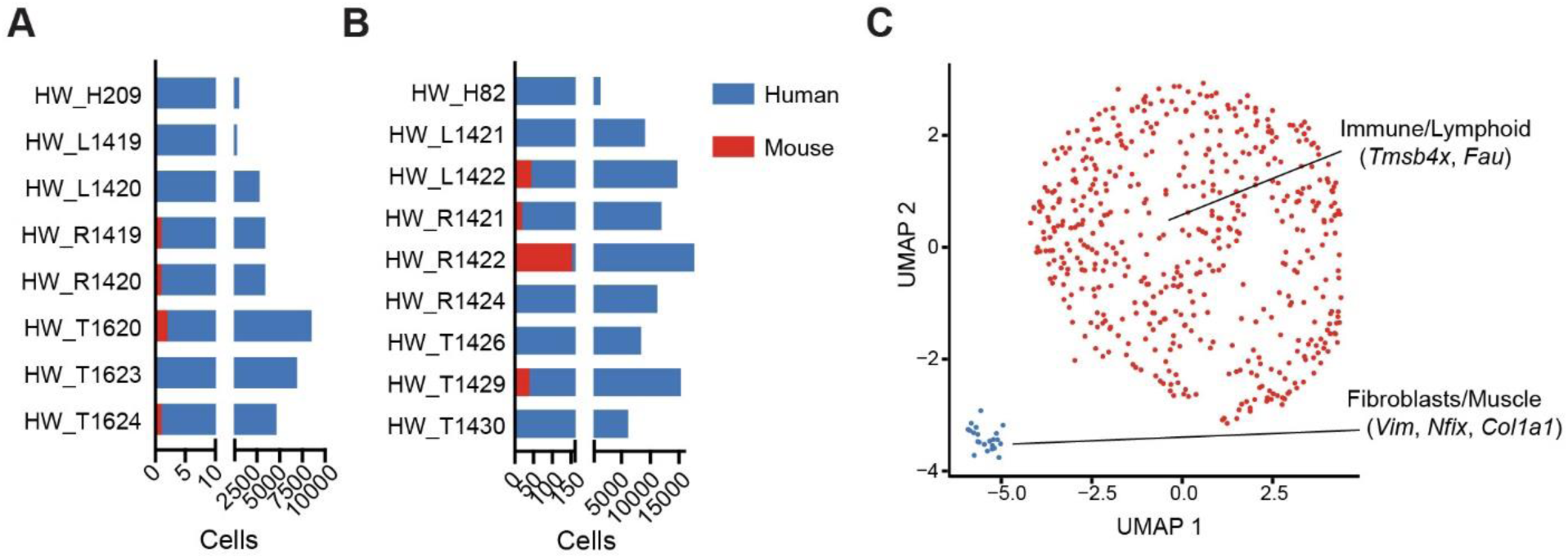
*In silico* removal and characterization of isolation of murine cells. Number of cell counts identified in each sample that maps to the human or mouse genome from NCI-H209 (**A**) and NCI-H82 cells (**B**). **C,** UMAP plot of the isolated murine cells from all samples cluster into two groups which are derived from either immune or fibroblast microenvironmental cells.

**Supplementary Figure S9.**
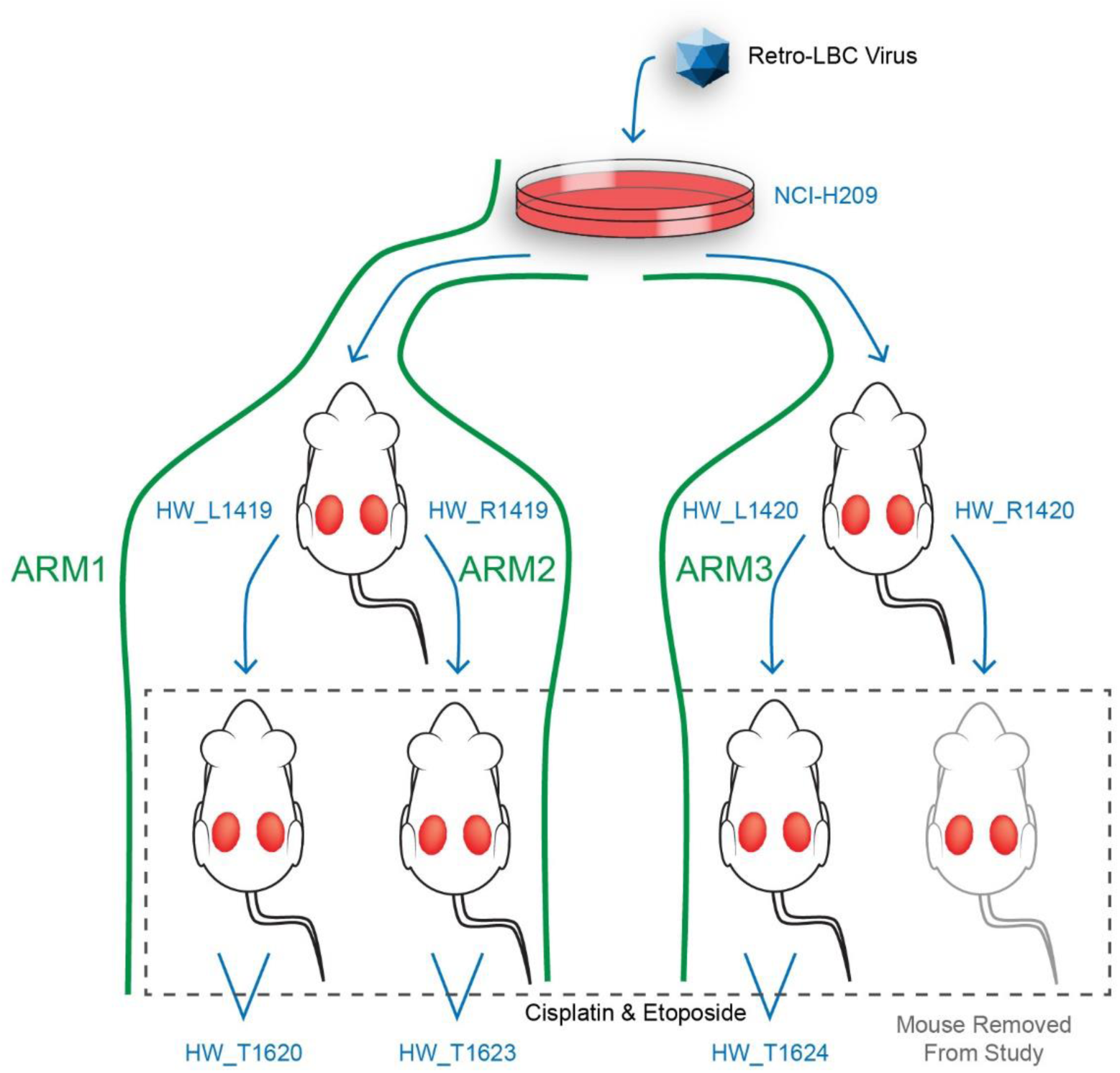
Graphic displaying the sample makeup of the three NCI-H209 sample arms. 2° transplant of HW_R1420 succumbed to condition unrelated to study before xenografts were palpable or chemotherapy treatment.

**Supplementary Figure S10.**
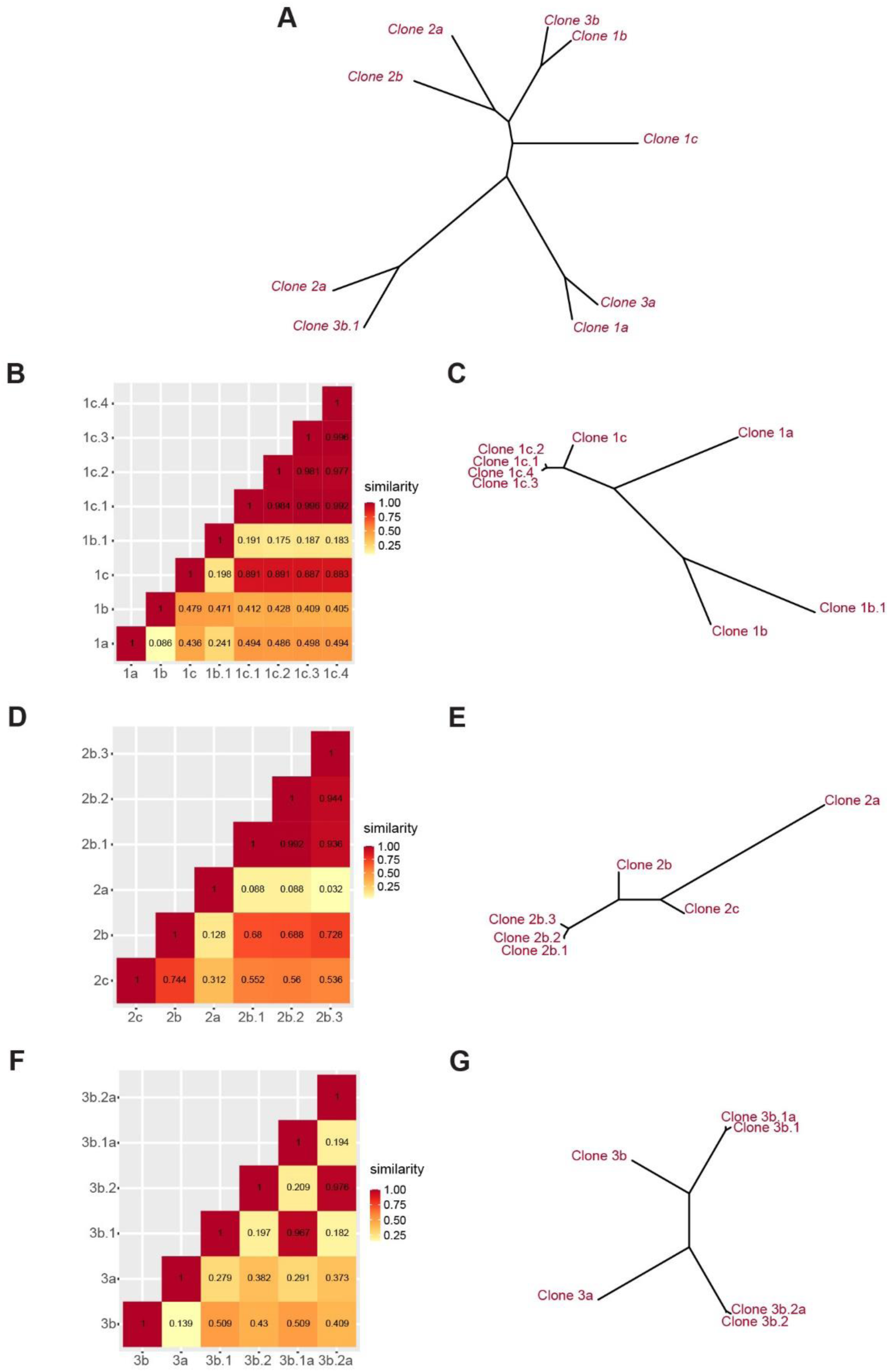
Similarity analysis to determine subclonal relationships. **A,** Neighbor-joining tree diagram showing relatedness of NCH-H209 subclone commonality between the three experimental arms. Similarity matrices (**B**, **D**, **F**) and evolutionary trees (**C**, **E**, **G**) for Arm 1 (**B**, **C**), Arm 2 (**D**, **E**), and Arm 3 (**F**, **G**).

**Supplementary Figure S11.**
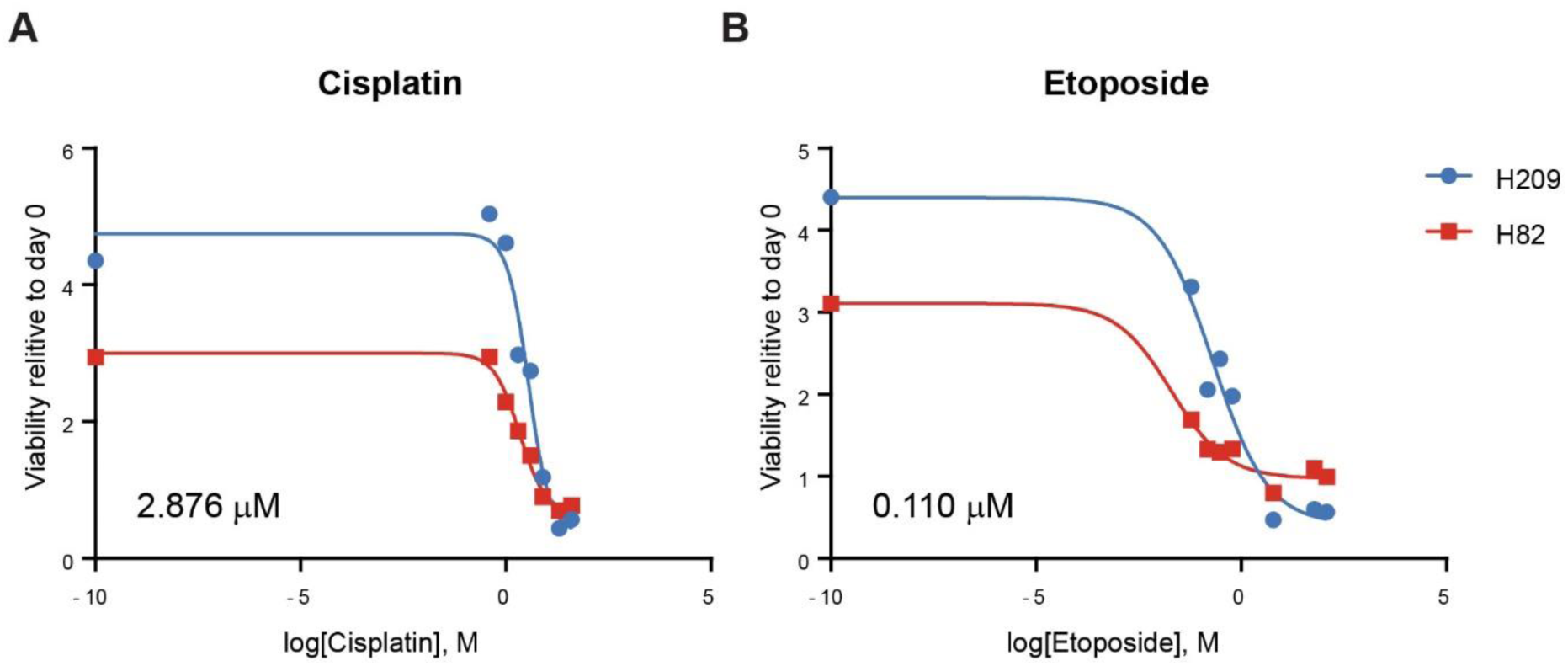
Determining effective concentrations for treatment of SCLC cell lines with cisplatin and etoposide. IC_50_ curves for the NCI-H20 9and NCI-H82 cells lines determined by an alamar blue assay for both cisplatin (**A**) or etoposide (**B**).

## Summary of additional Supplemental Tables

| **File** | **Description** | **File Type** |
| --- | --- | --- |
| Supplemental Table S3 | Gene module membership and expression of module 5 (related to Figure 1g) | Excel Spreadsheet (*.xlsx) |
| Supplemental Table S4 | Differential gene expression across NCI-H209 and NCI-H82 cluster (related to Figure 4) | Excel Spreadsheet (*.xlsx) |
| Supplemental Table S5 | Normalized gene expression across NCI-H209 subclones (related to Figure 4) | Excel Spreadsheet (*.xlsx) |
| Supplemental Table S6 | Normalized differentially expressed genes between subclones (related to Figure 5a) | Excel Spreadsheet (*.xlsx) |
| Supplemental Table S7 | Normalized CTA expression from NCI-H209 cell, xenografts, and chemo-treated xenografts (related to Figure 5f) | Excel Spreadsheet (*.xlsx) |
| Supplemental Table S8 | Oligos and Primer Sequences | Excel Spreadsheet (*.xlsx) |

